# Predictions of biodiversity are improved by integrating trait-based competition with abiotic filtering

**DOI:** 10.1101/2021.07.12.448750

**Authors:** Loïc Chalmandrier, Daniel B. Stouffer, Adam S. T. Purcell, William G. Lee, Andrew J. Tanentzap, Daniel C. Laughlin

## Abstract

All organisms must simultaneously tolerate the environment and access limiting resources if they are to persist. Otherwise they go extinct. Approaches to understanding environmental tolerance and resource competition have generally been developed independently. Consequently, integrating the factors that determine abiotic tolerance with those that affect competitive interactions to model species abundances and community structure remains an unresolved challenge. This is likely the reason why current models of community assembly do not accurately predict species abundances and dynamics. Here, we introduce a new synthetic framework that models both abiotic tolerance and biotic competition by using functional traits, which are phenotypic attributes that influence organism fitness. First, our framework estimates species carrying capacities that vary along abiotic gradients based on whether the phenotype tolerates the local environment. Second, it estimates pairwise competitive interactions as a function of multidimensional trait differences between species and determines which trait combinations produce the most competitive phenotypes. We demonstrate that our combined approach more than doubles the explained variance of species covers in a wetland community compared to the model of abiotic tolerances alone. Trait-based integration of competitive interactions and abiotic filtering improves our ability to predict species abundances across space, bringing us closer to more accurate predictions of biodiversity structure in a changing world.

## Introduction

Predicting species abundances is a major focus of community ecology (McGill et al. 2006). In recent decades, trait-based ecology has proposed that species morphological, physiological or phenological features determine how abiotic filtering and species interactions affect local community structure (Violle et al. 2007; Kraft et al. 2015b). However, trait-based analyses of communities often focus on functional diversity (Spasojevic et al. 2014; Chalmandrier et al. 2017) and few explicitly model species abundances (Zakharova et al. 2019).

Trait-based models of abiotic filtering that predict species abundances (Shipley 2010; Laughlin et al. 2012) assume that there are optimum trait values within a given environment, and species able to attain these trait values will be more likely occur in that environment (Kraft et al. 2015b). The most significant limitation of these models is that they fail to incorporate biotic interactions. In contrast, theoretical models of species interactions have a long and storied history in ecology (Lotka-Volterra 1925; Chesson 2000), and have been used to understand the foundational conditions for coexistence among competing species. For species to coexist stably, niche differences among species must be greater than differences in competitive ability (Chesson 2000; Adler et al. 2007) and recent work suggests that those differences can be linked to functional traits (Kraft et al. 2015a).

Three primary obstacles have prevented the mathematical integration of models of abiotic filtering and models of species interactions. First, they lack a common numerical currency through which they could be linked. Trait-based models of abiotic filtering yield probabilities that a species occurs in an environment given its traits, whereas models of species interactions describe dynamics of populations over time given growth rates, carrying capacities, and pairwise interaction coefficients (Lotka-Volterra 1925; Chesson 2000). Second, the complexity of estimating pairwise interactions increases exponentially with the number of species in the community, and there has been no obvious method for estimating interaction coefficients without implementing laborious competition experiments (Kraft et al. 2015a). Finally, there have been no adequate tools to model classical community ecology sampling schemes. For instance, plant abundance is often visually assessed through percent cover classes that do not necessarily fit well with existing statistical frameworks. Recently, authors have formalized the use of beta distributions to adequately model these sampling schemes (Damgaard & Irvine 2019), but they have yet to be implemented in biodiversity modeling.

Here we present a new synthetic framework that overcomes these three obstacles. This framework, which we call Banquo, integrates Traitspace, a trait-based model of abiotic filtering (Laughlin et al. 2012), with a Lotka-Volterra competition model. First, we assume that the probability that a species occurs in an environment given its traits is proportional to its local carrying capacity, *i.e*., the maximum population size that a species can reach given local resources and abiotic conditions in the absence of competition (MacArthur & Levins 1967). Second, we assume pairwise interaction coefficients are a function of observed trait differences between species, thereby substantially reducing the number of parameters needed to estimate pairwise interaction coefficients (Chalmandrier et al. 2021). Drawing inspiration from coexistence theory (Chesson 2000; Adler et al. 2007), the parameterization of this function allows for competitive outcomes to be affected to differing degrees by both niche partitioning (*i.e*., strong competitive interference among functionally similar species) and competitive hierarchies (*i.e*., species have strong competitive impacts on species with inferior trait values). Third, we use the recent methodological developments of Irvine et al. (2019) to link the output of our framework to observed plant species abundances that were estimated through cover classes.

We illustrate our framework by modeling plant species abundances along a flooding gradient in an ephemeral wetland (Purcell et al. 2019). After presenting our framework, we calibrated sixteen assembly models that include abiotic filtering and/or biotic filtering tested on different sets of functional traits. Then, we compared the statistical performance of these sixteen assembly models. Finally, we analyzed how the parameterization and output of the calibrated models inform our knowledge about the assembly of wetland plant communities.

## Methods

### The framework

#### Step 1 – Estimating species carrying capacities along environmental gradients

We started with the Traitspace framework to model species’ probabilities of occurrence along the flooding gradient (Laughlin et al. 2012). Traitspace characterizes the size and shape of the environmental filter based on a multivariate linear model with a vector of individual plant traits (*T*) as the response and a vector of environmental gradients (*E*) as the predictors, i.e. the function *T* = *f*(*E_k_*). Traitspace uses this linear model to estimate the conditional distributions of traits *T* given the environmental conditions in site *k* (*P*(*T|E_k_*)). Second, it uses the intraspecific trait distribution of each species across sites, i.e. the conditional distributions of traits given species identity (*P*(*T|S_i_*)). The posterior distribution of species presence *S_ik_* of species *i* in site *k* is conditioned on both the trait state *T* and the environmental conditions *E_k_*. *P*(*S_ik_|T,E_k_*) is computed using Bayes theorem:

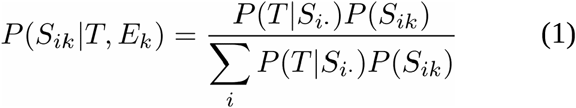

The desired posterior is computed by integrating with respect to traits to obtain the probability of occurrence of a species given the environmental conditions:

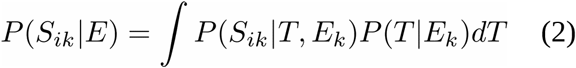

In practice, we use Monte Carlo integration to estimate the average probability of presence of each species in each site by randomly sampling 500 trait values per site based on the estimated trait-environment relationship (*T* = *f*(*E*)) and then averaging the probability distribution for each site and each species. In the end, we obtained a site-by-species probability table.

We then assumed that the carrying capacity (in percent cover) *K_ik_* of species *i* in a site *k* can be estimated from its probability of presence in that site using a increasing log-log function:

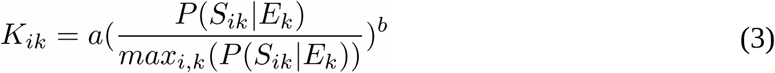

with *a* ∈ [0,1], 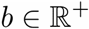.

We standardized the probability value *P*(*S_ij_|E_j_*) by the maximum value across all species *i* and all sites *j* to ensure that carrying capacities *K_ij_* are all set between 0 and 1 (as a percent cover variable).

#### Step 2 – Modeling the biotic filter: estimation of trait-mediated plant competitive interactions

**Formulation of the interaction matrix –** Here we assume that the interaction coefficient *α_ik_* that measures the competitive impact of species *j* on species *i* can be estimated as a function of difference in traits. We test a formulation of *α_ij_* as a function of the empirical trait of value *t_i_* of species *i* and *t_j_* of species *j*:

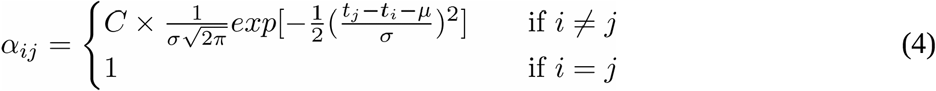

with 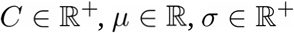.

Interspecific coefficients followed a modified Gaussian function of trait differences where *μ* is the peak position of the Gaussian, *σ* is its width, and *C* controls the amplitude of interspecific coefficients relative to intraspecific coefficients. Species intraspecific coefficients were fixed to 1. For a small values of the ratio 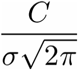, the matrix of interaction coefficients can be approximated by the identity matrix (***α*** = ***I***) and estimated species covers simplify to the vector of carrying capacities. For large values of *σ* (*σ*→∞), interspecific coefficients are all equal to 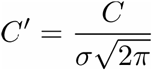 and represent a situation where interspecific interactions among species are constant and do not depend on species traits.

The formulation of equation 4 is that it can be directly related to either competitive hierarchies or niche partitioning (Chesson 2000; Adler et al. 2007). The value of the parameter *μ* defines if the studied trait relates more to niche partitioning among species, hierarchical competition, or a mixture of the two. Specifically, for *μ* close to 0, pairwise interaction coefficients are high for small trait differences and low for large trait differences, indicating a predominance of niche partitioning among species (Supplementary figure 1-A). For a low value of *μ*, the left hand part of the bell-shaped curve falls outside of the range of observed trait differences. Thus the curve approaches a monotonically decreasing function that indicates a predominance of competitive hierarchy: species with a large trait value are competitively superior over species with small trait values (Supplementary figure 1-B). Conversely, a high value of *μ* returns a monotonically increasing function that indicates a predominance of hierarchical competition with species with a small trait value being competitively superior over species with large trait values. Intermediate situations (moderately large or moderately small values of *μ*) indicate a mixture of niche partitioning and hierarchical competition (Supplementary figure 1-C): niche partitioning is predominant among species with large trait differences, but among species with small trait differences, competition is not symmetric (as in a case of “pure” niche partitioning”) and hierarchical competition is the predominant coexistence process.

Finally, our formulation of interaction coefficients can be extended to multiple trait dimensions using a modified multivariate Gaussian function. In this study, we used a maximum of two trait dimensions in which case interaction coefficients were formulated as follows:

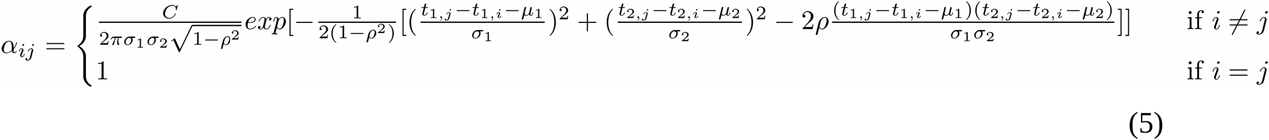

with 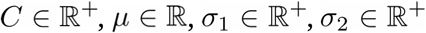, |*ρ*| < 1.

This equation describes a two-dimensional symmetric Gaussian function of trait differences of peak position (*μ_1_, μ_2_*) and of widths *σ_1_* and *σ_2_* across the first and second dimensions. Properties and interpretations of the parameters are similar to their uni-dimensional counterparts. The two-trait formulation includes an additional coefficient *ρ* between the two trait difference dimensions that determine if trait differences independently contribute to the pairwise interaction coefficients (*ρ* = 0) or if they interact (0 < |*ρ*| < 1).

#### Step 3 - Integrating the abiotic and biotic filter with Lotka-Volterra models

We assumed that species’ dynamics could be modeled though a Lotka-Volterra competition model:

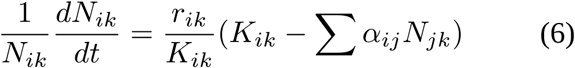

where *N_ik_* and *r_ik_* are, respectively, the percent cover and the intrinsic growth rate of species *i* in site *k*.

Within this model, the vector of all strictly positive species covers at equilibrium 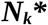 satisfies the equation:

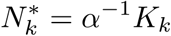

where ***K_k_*** = {*K_ik_*} is the vector of species carrying capacities, and ***α*** = {*α_ij_*} is the matrix of percapita effects estimated as described above.

For a given set of parameters, the interaction matrix α was estimated, the Moore-Penrose inverse of ***α*** was computed, and multiplied to each site’s vector of species carrying capacities estimated from the Traitspace model. Species local cover estimated in this way can be negative, reflecting that this equilibrium state is not feasible. To find a feasible equilibrium, for each vector of equilibrium species covers, we sequentially set to 0 the species with the most negative cover and re-estimated the equilibrium state. This procedure was repeated until finding an equilibrium state where all remaining species covers were positive.

### A test of the framework

We tested our framework on a dataset of an ephemeral wetland in New Zealand (latitude 44.374143°S, longitude 169.890052°E). In that ecosystem, plant community structure vary along a continuous flooding gradient. In this test, we assumed that plant community assembly is determined by the filtering of three functional traits by flooding duration that filters three functional traits (root porosity, height and SLA) and above-ground competition determined by height and SLA.

Analyses of the dataset are available in previous studies (Tanentzap et al. 2014; Tanentzap & Lee 2017; Purcell et al. 2019). Detailed methods about data collection are available in the supplementary materials. We analyzed the vegetation structure with a subset of the complete dataset (see Supplementary materials): 67 quadrats 25 × 25 cm in size set along four transects that run from the lowest point of the basin and advancing upslope to the kettlehole margin. Foliar cover was estimated for each species using the following cover estimates: 0.5%, 1%, 2%, 3%, 4%, 5%, 10%, 15%, 20%, 30%, 40%,…, 100%. We restricted the analysis to the 15 most abundant species in the study area for which we sampled traits on at least 20 individuals. These species collectively represent at least 80% of the total of plant cover in each quadrat (Pakeman & Quested 2007).

Root porosity, as a percentage variable, was logit-transformed. Height and SLA trait values were log-transformed prior to the analysis to approach a normal distribution. We modeled the relationship between root porosity, SLA, height and the flooding gradient and weighted trait observations by species cover.

The intraspecific trait distribution of each species was modeled using a multivariate normal distribution (R-function *mclust::dens Scrucca et al. 2016*). We then modeled the probability of occurrence of each species in each site given the local duration of flooding using the Traitspace framework described earlier.

To calibrate the interaction matrices, we used species maximum height along the gradient (calculated as the 95% quantile of each species height values) and species average SLA. Maximum height and SLA were moderately correlated (*r* = −0.42, *t* = −1.65, *df* = 13, *p* = 0.12). To avoid using correlated functional traits to estimate the two-traits interaction matrices, we first computed a PCA on the species by trait matrix containing species maximum height and average SLA. We then used species scores along these two PCA trait axes to calibrate the pairwise interaction matrix. As we used all the PCA dimensions, this step does not compromise the amount of trait variation used to estimate the two-trait interaction matrix. Practice showed us that, compared to using correlated (but tangible) functional traits, this extra step facilitates and speeds the convergence of the model calibration algorithm described below. However, we related pairwise interaction coefficients to the observed species functional traits values, rather than to the PCA trait axes, to facilitate the ecological interpretation of our results.

Using the Banquo framework, we tested a total of sixteen assembly models. All sixteen assembly models aim to solve the following equation to estimate the matrix of species cover *N*\* at equilibrium:

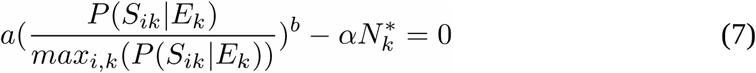

(Model 1) One null model without any assembly processes: species probability of presence given local abiotic conditions were assumed to be equally abundant in every site (*b* = 0) and there is no interspecific competition (the interaction matrix ***α*** is equal to the identity matrix ***I***).

(Model 2) One abiotic filtering model: species probability of presence are estimated by the Traitspace framework (b > 0) and there is no interspecific competition (***α*** = ***I***).

(Models 3-7) Five biotic filtering models that include no abiotic filtering: species probability of presence were assumed to be equal across species and in every site (*b* = 0) but species cover is determined by interspecific competitive interactions (***α*** ≠ ***I***) that could depend on 3) no traits (*σ*→∞), 4) plant height, 5) SLA, 6) both height and SLA without the interaction parameter *ρ*, or 7) both height and SLA with the interaction parameter ρ.

(Models 8-12) Five abiotic and biotic filtering models: species probability of presence are estimated by the Traitspace framework (*b* > 0) and species cover is also impacted by interspecific competitive interactions (***α*** ≠ ***I***) that could depend on 8) no traits (*σ*→∞), 9) plant height, 10) SLA, 11) both height and SLA without the interaction parameter *ρ*, or 12) both height and SLA with the interaction parameter *ρ*.

We summarize the characteristics of each assembly model and their parameters in Table 1.

**Table 1.** Comparison of the assembly models. Each model posterior is described by the Deviance Information criterion (DIC), Nagelkerke’s pseudo R^2^, the area under the curve (AUC) and Gelman’s multivariate convergence criterion (MPSRF). The assembly models are ranked by decreasing median pseudo R^2^.

| Assembly model category | Traits used to calibrate interactions ($\alpha_{ik}$) | Fixed parameters | Parameters to estimate | MPSRF | DIC | Pseudo $R^2$ | AUC |
| --- | --- | --- | --- | --- | --- | --- | --- |
| Abiotic & Biotic model | SLA | / | $a, b, C, \mu, \sigma, \varphi$ | 1.002 | 2291 | [0.182, 0.196] | 0,773 |
| Abiotic & Biotic model | Height + SLA (with int.) | / | $a, b, C, \mu_1, \mu_2, \sigma_1, \sigma_2, \rho, \varphi$ | 1.004 | 2295 | [0.177, 0.194] | 0,773 |
| Abiotic & Biotic model | Height + SLA (no int.) | / | $a, b, C, \mu_1, \mu_2, \sigma_1, \sigma_2, \varphi$ | 1.018 | 2308,8 | [0.171, 0.188] | 0,764 |
| Abiotic & Biotic model | Height | / | $a, b, C, \mu, \sigma, \varphi$ | 1.000 | 2371,8 | [0.115, 0.13] | 0,689 |
| Abiotic model | No interactions | $\alpha = I$ | $a, b, \varphi$ | 1.000 | 2413,8 | [0.075, 0.085] | 0,671 |
| Abiotic & Biotic model | No traits | $\sigma \rightarrow \infty$ | $a, b, C', \varphi$ | 1.001 | 2412 | [0.071, 0.082] | 0,669 |
| Biotic model | Height + SLA (with int.) | $b = 0$ | $a, C, \mu_1, \mu_2, \sigma_1, \sigma_2, \rho, \varphi$ | 1.811 | / | [0.056, 0.088] | / |
| Biotic model | Height + SLA (no int.) | $b = 0$ | $a, C, \mu_1, \mu_2, \sigma_1, \sigma_2, \varphi$ | 1.021 | 2415,2 | [0.038, 0.067] | 0,657 |
| Biotic model | SLA | $b = 0$ | $a, C, \mu, \sigma, \varphi$ | 1.006 | 2420,7 | [0.03, 0.043] | 0,631 |
| Biotic model | Height | $b = 0$ | $a, C, \mu, \sigma, \varphi$ | 1.025 | 2456 | [0.027, 0.042] | 0,574 |
| Biotic model | No traits | $b = 0, \sigma \rightarrow \infty$ | $a, C', \varphi$ | 1.000 | 2491,8 | [0, 0.001] | 0,448 |
| Null model | No interactions | $\alpha = I, b = 0$ | $a, \varphi$ | 1.000 | 2491,8 | [0, 0.001] | 0,5 |

#### Calibration and comparison

We used the likelihood function proposed by Irvine et al. (2019) given that observed species covers were recorded as percent cover classes. Briefly, the likelihood function links the ordinal observations of plant cover to a latent beta distribution of mean *N_ij_* (in our case estimated by the assembly models) and uncertainty parameter *φ*, that can be interpreted as a measure of plant spatial aggregation (Damgaard & Irvine 2019). One drawback of using the beta distribution is that it cannot model zero percent covers. To circumvent that issue, we added a small offset (0.05 %) to zero percent cover values, as suggested by Irvine et al. (2019). Note that this corresponds to moving unobserved species up to the next highest cover class, that only included a single cover values (0.1% of the total number of cover values).

Depending on the assembly models, there were two (null model) to nine parameters (abiotic + biotic model with height and SLA with the interaction parameter *ρ*) to estimate. We set regularizing priors on all parameters (Banner et al. 2020): we avoid making *a priori* assumptions about the nature of the relationship between traits, carrying capacities, pairwise interactions and species cover but we limited the extent of the parameter space that was uninformative. Details about the prior functions and their hyper-parameterization are available in the supplementary materials.

We used a Differential-Evolution Markov-Chain Monte Carlo algorithm (DEzs MCMC in the R-package BayesianTools (Hartig et al. 2017) to estimate the posterior distributions of the parameters. For each model, we ran four chains for 6 x 10^5^ steps. Convergence was assessed through Gelman’s multivariate convergence criterion (MPSRF, Gelman et al. 2014).

#### Assembly model comparison –

We compared the fits of the calibrated models using two metrics: the Deviance Information Criterion (DIC, Gelman et al. 2014) and Nagelkerke’s pseudo R^2^ metric (Nagelkerke 1991) which lends itself well to models that use a beta distribution and gives an indication of the variance they explain (Nakagawa & Schielzeth 2013). Nagelkerke’s pseudo R^2^ was calculated from the ratio of a model’s posterior likelihood and the likelihood of the null model (see above). Furthermore, we evaluate the ability of the models to predict species presence/absence by evaluating receiver operating characteristic curve (ROC) and the area under the curve (AUC) scores of each assembly model of the predicted species-site matrix using the R-package pROC (Robin et al. 2011).

#### Code availability

The R-scripts and data to run the analysis are available at https://github.com/LoicChr/Banquo

## Results

### Relationship between flooding and functional traits

Root porosity increased significantly with flooding duration (*t* = 7.714, *df* = 224, *P* < 0.0001. Adjusted R^2^ = 0.206, Figure 1). Plant height decreased (*t* = −3.91; *df* = 224, *P* = 0.0001) and specific leaf area increased with flooding duration (*t* = −3.10; *df* = 224, *P* = 0.002) but these relationships explained only a negligible portion of trait variation along the flooding duration gradient (Height adjusted *R^2^*: 0.036; SLA adjusted *R^2^*: 0.060).

**Figure 1.**
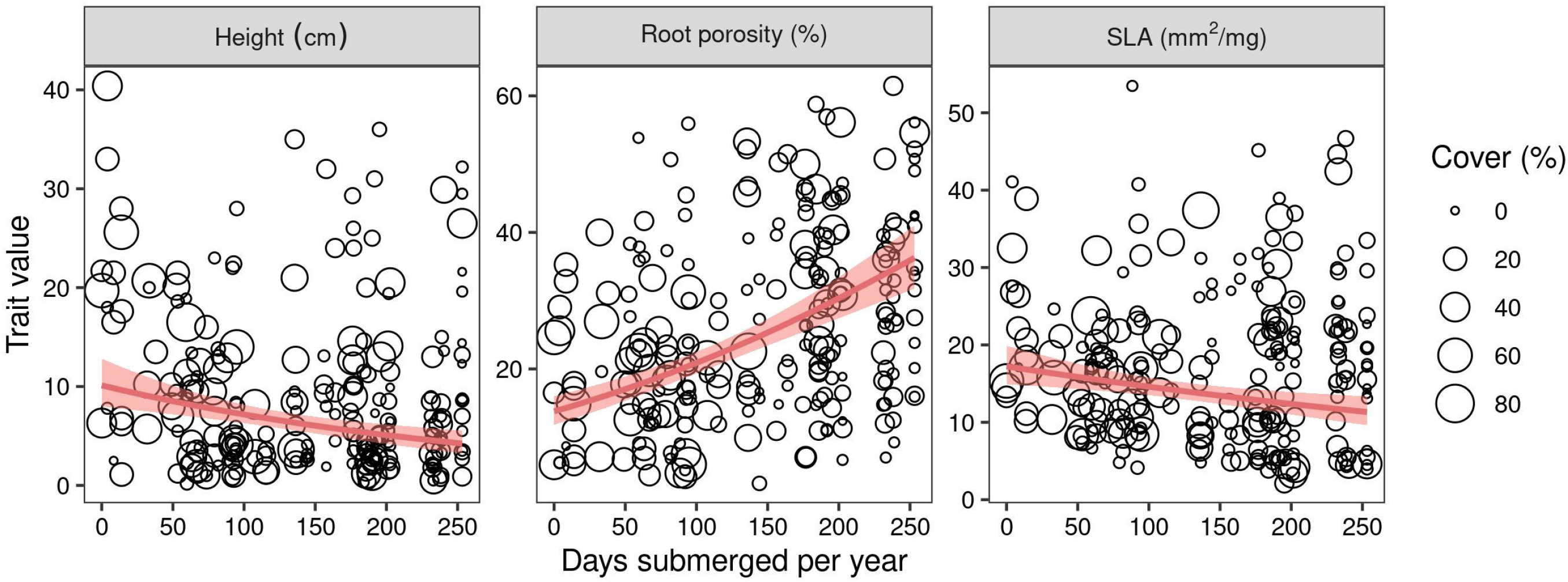
Relationship between the duration of flooding and plant traits: root porosity (a), specific leaf area (b) and vegetative height (c). Data point size is proportional to plant cover. The line indicates the modeled relationship used in the Traitspace framework. The three linear models were all statistically significant (Root porosity, adjusted R^2^ = 0.206, p < 1 × 10^-5^; Specific leaf area, adjusted R^2^ = 0.037, p = 0.0021; Vegetative height, adjusted R^2^ = 0.060, p =0.0001).

### Model comparison

All assembly models but one converged (Gelman’s multivariate convergence criterion inferior to 1.1). Regardless of the performance statistic, there was a clear hierarchy across the assembly models (Table 1). The biotic models without abiotic filtering performed the worst (DIC: [2415.2, 2491.8], median pseudo R^2^: [0.001, 0.088], AUC: [0.44, 0.68]). The biotic model calibrated with both height and SLA (with interaction term) had a convergence criterion of 1.81; we thus could not calculate its DIC and AUC. The associated pseudo R^2^ was however low across its posterior distribution: [0.056, 0.088]. The abiotic model without biotic interactions performed better (DIC: 2413, median pseudo R^2^: [0.075, 0.085], AUC: 0.67).

The models that included both abiotic filtering and biotic interactions performed the best both in explained plant cover variance (pseudo R^2^: [0.082, 0.196]; DIC: [2291.0, 2412.0]) and species presence/absence (AUC: [0.67, 0.77]). The model that assumed fixed pairwise interaction coefficients among competitive species was the worst performing of all (pseudo R^2^: 0.081; DIC: 2412.0, AUC = 0.67), while the SLA-based interaction models were the best performing. Among the latter, the model that calibrated biotic interactions using specific leaf area was the best fitting (DIC: 2291.0, pseudo R^2^: 0.196, AUC: 0.77).

### Calibrated pairwise interaction matrices

Among the assembly models that included both abiotic filtering and a pairwise interaction matrix calibrated with functional traits, the pairwise interaction matrix calibrated with SLA was the best supported by the data. It indicated a predominance of niche partitioning. There was a slight but non-significant hierarchical competition effect (μ: 95% IQ [-0.43, 0.08], Figure 2C). This showed that the modest hierarchical competition among pairs of species conferred an advantage to species with the largest SLA.

**Figure 2.**
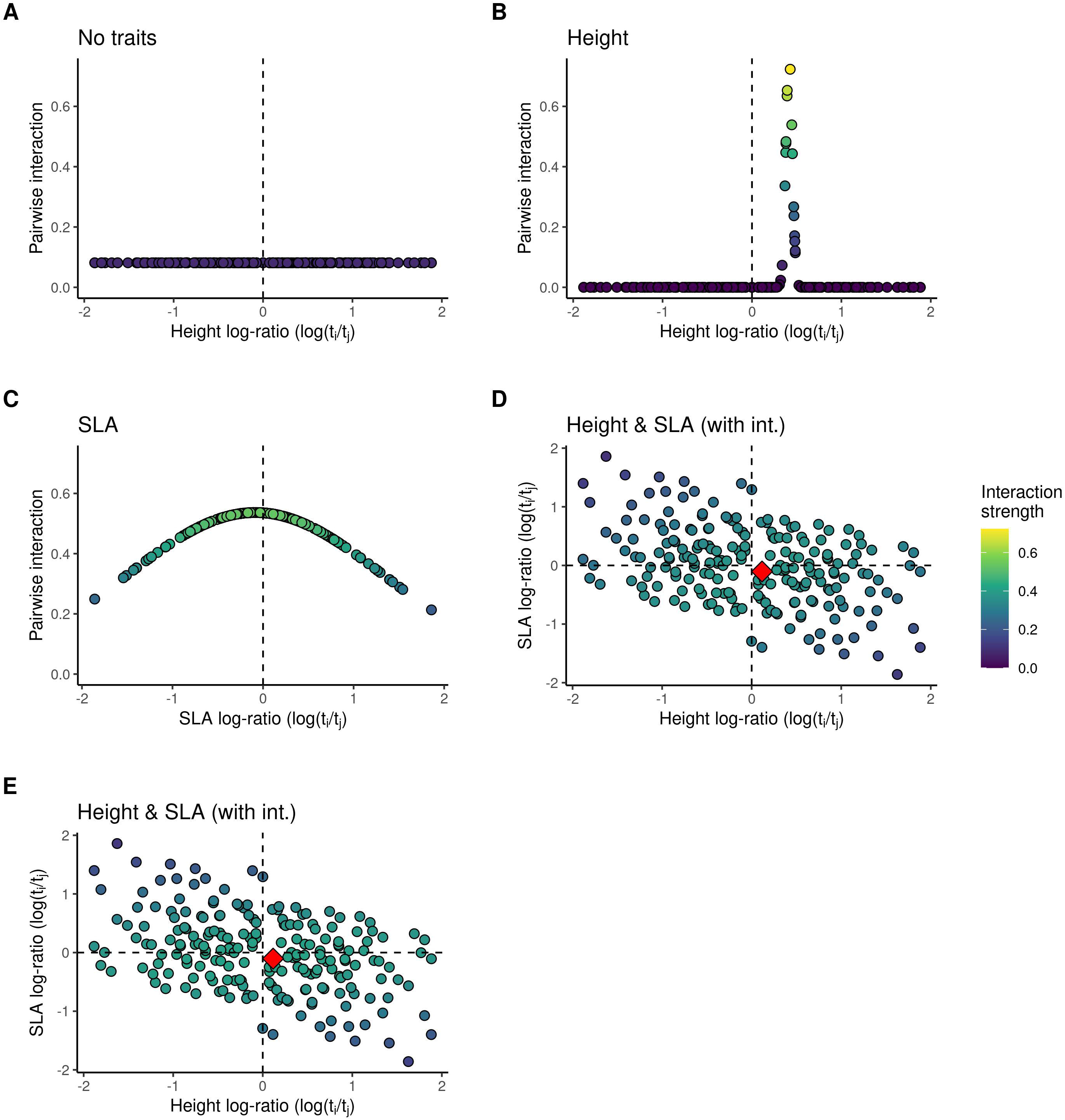
Calibrated pairwise interaction coefficients of the abiotic and biotic assembly models. The graphic represents interspecific *α_ij_* (competitive impact of species *j* on species *i*) as a function of the height-only (a), SLA-only (b), or both (c,d) differences of species *i* and species *j*. The blue scale represents the absolute value of the pairwise coefficient. To facilitate the interpretation of the two-traits plots (c-d), we indicated the position of the largest pairwise interaction coefficient value with an orange diamond.

All three pairwise interaction matrices calibrated with SLA were similar regardless of the inclusion of height or the interaction parameter *ρ*. At the median of their respective posterior distribution, the pairwise interspecific competition coefficients were strongly correlated (*r* = 0.82 between the SLA-calibrated matrix and the Height + SLA calibrated matrix without interaction; *r* = 0.98 between the SLA-calibrated matrix and the Height + SLA calibrated matrix with interaction).

### Comparison between the abiotic model and the abiotic and biotic assembly models

We compared the abiotic model to the best assembly model (i.e. abiotic and biotic with pairwise interactions calibrated with Height and SLA without interaction). The abiotic model tends not to predict species absences well. The distribution of cover values was thus approximately normal around a median value of 3.30% (Supplementary Figure 5). Consequently, species presence along the flooding gradient was often overestimated with numerous species being predicted to be present in sites where they were not observed (e.g. see *Epilobium angustum*, Figure 3). In contrast, when biotic interactions are included, the assembly model tends to predict more absences and less even percent cover values among species (Figure 3, Supplementary Figure 5).

**Figure 3.**
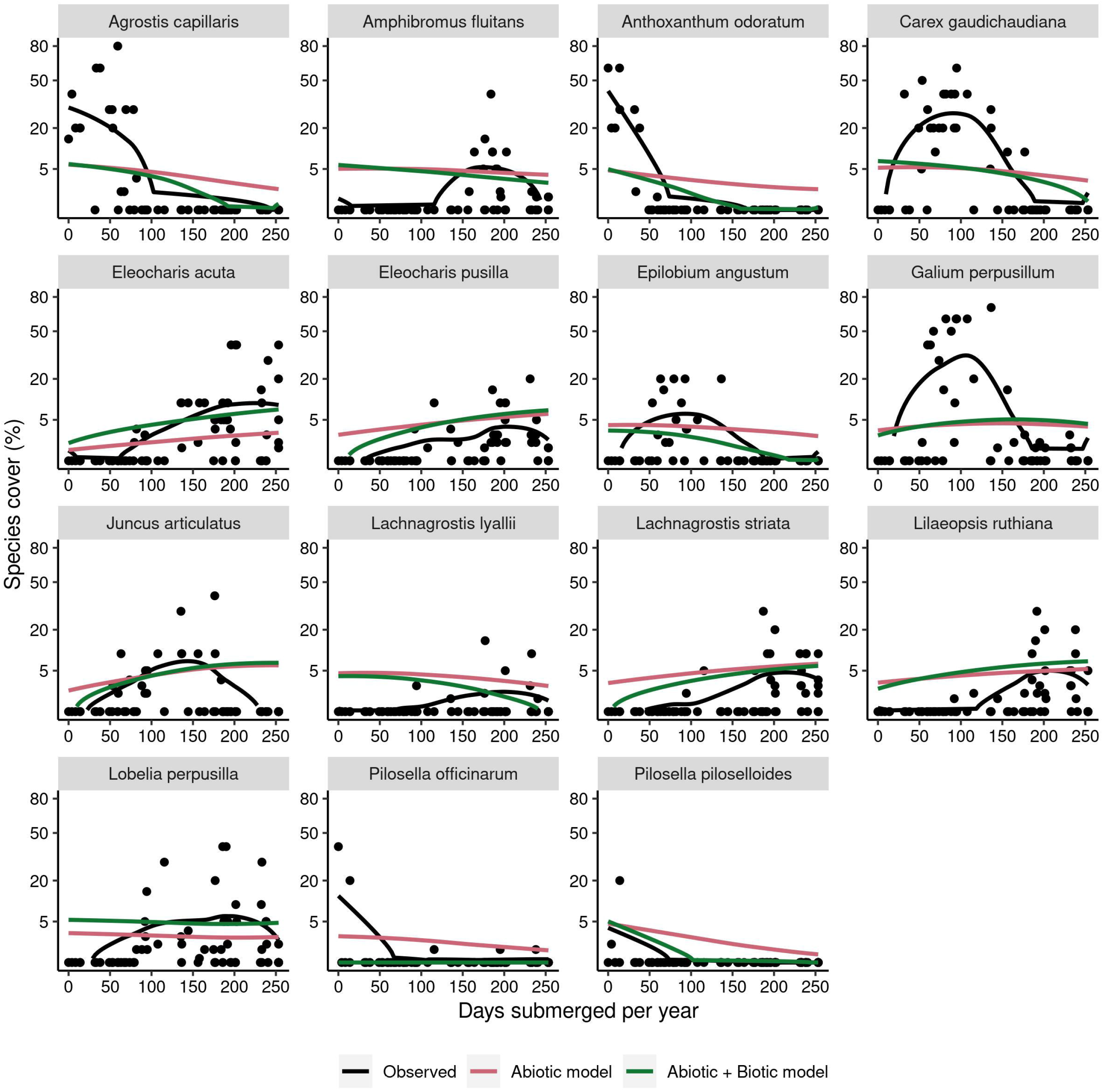
Comparison of observed and modeled species cover along the flooding gradient. Red response curves are fitted on the species cover as predicted by the abiotic model. Green response curves are fitted on the species cover as predicted by the best abiotic and biotic model (see Table 1). Black nonlinear response curves are fitted directly to the observed cover for each species. All curves are fitted using a loess function.

## Discussion

Predictive models of community assembly have focused on incorporating abiotic filters and have generally ignored biotic interactions. Here we show that trait-based assembly rules can be used to directly model species abundances in communities by simultaneously accounting for both abiotic filtering and competitive interactions (Keddy 1992, 2001). There are two major implications of this study. First, we introduced a trait-based formulation of pairwise competitive interactions that allowed us to calibrate 210 interaction coefficients from observational data with no more than eight parameters. This new approach substantially improves our ability to infer interaction matrices with little additional complexity (Cabral et al. 2017; Chalmandrier et al. 2021). Second, the inclusion of competitive interactions among species improved predictions of local plant cover, which bolsters the argument that the modeling of species distribution must include both abiotic tolerances and species interactions (Alexander et al. 2015; Evans et al. 2016).

The core feature of the Banquo model is the formulation and calibration of pairwise competition coefficients among species. We proposed a new flexible formulation of competitive pairwise interactions as a function of trait differences. Compared to estimating pairwise interaction coefficients individually, this considerably reduces the number of parameters to estimate (Zakharova et al. 2019; Chalmandrier et al. 2021). That formulation was directly inspired by, and thus constrained by, the principles of coexistence theory and how it has informed the study of functional diversity patterns (Chesson 2000; HilleRisLambers et al. 2012).

Traditionally, functional diversity pattern studies have assumed that niche partitioning was the main competition mechanism behind community assembly (MacArthur & Levins 1967). In that framework, niche partitioning would promote the coexistence of functionally dissimilar species and oppose itself to environmental filtering that promotes the coexistence of functionally similar species. In recent years, this framework has been criticized (Kraft et al. 2015b; Münkemüller et al. 2020) as coexistence theory posits that competition can also promote the coexistence of functionally similar species through hierarchical competition. Our framework has the benefit of not assuming niche partitioning or hierarchical competition as the main trait-based competitive mechanism among species, but rather permits the fit of a mixture of the two processes. Our empirical example illustrates that the pairwise interaction matrices of the assembly models were consistent with strong niche partitioning among species but with a small yet significant degree of hierarchical competition among species with small trait differences. Our modeling of pairwise interactions aims to provide a general and flexible relationship between competition strength among plants and trait differences rooted in coexistence theory. However, it is essentially phenomenological and does not explicitly model the mechanisms behind plant-plant competition. Future developments may aim at formulating competition as an explicit function of species’ ability to consume local soil resources Letten et al. (2017) or intercept light (Falster et al. 2017). Beyond competition, a more complex modeling of plant biotic interactions could include facilitative interactions or acknowledge that the nature of species interactions can shift along environmental gradients (Maestre et al. 2009; Bimler et al. 2018).

Our trait-based modeling approach explicitly specifies classical assembly mechanisms (HilleRisLambers et al. 2012) and evaluates their ability to predict species local abundance using common modeling statistics. Our case study showed that including both abiotic filtering and biotic interactions led to a net improvement of the modeling of species local abundances and of community structure. One of the limitations of established trait-based models is that they tend to overestimate species occurrences (e.g. Merow et al. 2011). This drawback also affects other types of biodiversity models such as stacked species distribution models (Pottier et al. 2013) leading to inaccurate predictions of community structure at small spatial scales (Thuiller et al. 2015). This has long been interpreted as a consequence of not properly accounting for biotic interactions. Our study supports for this conjecture: including competitive interactions improves the modeling of species occurrences and further decreases the predicted diversity (α-diversity) and increases the predicted turnover (β-diversity) bringing them closer to the observed diversity values (Supplementary Figure 4). Our results follow the classical expectation that the realized niche of species is smaller than the fundamental niche because species interactions limit where species actually occur (MacArthur & Levins 1967). In more details, the assembly model that include only abiotic filtering predicts remarkably even species abundances (Figure 3, Supplementary figure 5), in contrast with the usual strong heterogeneity that characterizes species abundance distributions (McGill et al. 2007). The inclusion of biotic interactions predicts a more realistic distribution of species abundances within communities and produces more sparse species-by-site community matrices that exhibited a stronger hierarchy among species (Figure 3, Supplementary figure 5).

By applying our framework to the strong flooding gradient in a wetland ecosystem, we were able to get insights into the ecological mechanisms that determine wetland community structure and also identify our framework limitations. First, we found that a trait-based model of abiotic filtering (root porosity, SLA, and height) led to a modest improvement in model fit compared to the null model (Table 1, pseudo *R*^2^ 95% IQ [0.075, 0.085]). This suggested flooding filtered the species pool primarily by porous root tissue that enhances the ability of species to tolerate flooded and anoxic soil (Moor et al. 2017; Tanentzap & Lee 2017). When we added trait-based competitive interactions to this assembly model, we significantly improved the modeling of species covers and, consequently, of community structure. Using traits to estimate the interaction matrix further proved useful as the assembly model with a fixed pairwise interaction matrix was not well supported by the data. The “best” model was the model that calibrated biotic interactions with SLA (Table 1, pseudo *R*^2^ 95% IQ [0.182, 0.196]). This suggests that competitive interactions among plants in that ecosystem could be mediated through leaf economics (Violle et al. 2009; Tanentzap & Lee 2017). In contrast, the interaction matrix calibrated only with height was less supported by the data. This indicated that there was little competitive interference among pairs with dissimilar SLA values, likely because they partition resources and are thus able to coexist (Moor et al. 2017).

However, even the best assembly model explained a relatively modest portion of species abundances. This points both to the limitations of the available data and of our framework. Only root porosity was found to vary, and only moderately, along the flooding gradient (adjusted R^2^ = 0.21). Thus the modeled carrying capacities of species along the flooding did not vary as strongly as could be *a priori* expected (see Figure 3). The intraspecific variability of root porosity was important (32% of total root porosity variance was intraspecific) and may dampen our ability to use this trait to model species’ abiotic niche (Read et al. 2017). It is also possible that other unmeasured functional traits may be involved in the filtering of species along the flooding gradient (Moor et al. 2017).

## Conclusion

It has been argued that complex ecological processes can be modeled with limited data input by leveraging the generality of functional traits (McGill et al. 2006). In community ecology, functional traits are mainly used in diversity pattern analyses codified by assembly theory (Keddy 1992). Those analyses have numerous pitfalls: non-random functional diversity patterns can be interpreted in multiple ways thus rendering difficult a confident inference of community assembly rules (Kraft et al. 2015b; Cadotte & Tucker 2017; Münkemüller et al. 2020). In contrast, our approach specifies explicit assembly rules and model directly local species abundances. Ultimately, our framework provides a process-based approach to predict community structure and quantify its support. Such trait-based modelling opens a new general way to model natural communities and will improve our ability to understand and predict biodiversity structure and dynamics under global change.

## Acknowledgments

This research was supported by funding from both the University of Waikato and Manaaki Whenua - Landcare Research. John Payne of Landcare Research kindly provided the capacitance probe water level data. We call our new model, ‘Banquo’, in honor of Bill Shipley, who, in his foundational book (Shipley 2010) was the first to highlight Banquo’s desire to predict ‘which grain will grow and which will not’ in Shakespeare’s Macbeth. LC acknowledges a postdoctoral fellowship at the University of Wyoming and funding from the European Union’s Horizon 2020 research and innovation program under the Marie Skłodowska-Curie grant agreement No 840946 (Project “CLIMB”). DBS acknowledges the support of a Rutherford Discovery Fellowship and the Marsden Fund Council from New Zealand Government funding, both of which are managed by the Royal Society of New Zealand Te Apārangi (RDF-13-UOC-003 and 16-UOC-008), and an Erskine Grant from the University of Canterbury.

## Author Contributions

LC, DBS and DCL led the study. LC developed and ran the analyses and wrote the first draft. ASTP, WGL, AJT and DCL collected the data. All authors contributed to the writing of the paper.

## Notes

https://github.com/LoicChr/Banquo

### Competing Interest Statement

The authors have declared no competing interest.

