## Supplementary information for "Predictions of biodiversity are improved by integrating trait-based competition with abiotic filtering"

#### 1 Supplementary Methods

##### 1.1 Study site

The data were collected in an ephemeral wetland in a kettlehole in the Mackenzie Basin, South Island, New Zealand (latitude 44.374143°S, longitude 169.890052°E) at an elevation of 599 m. The kettlehole was formed by glacial settling and is nested gently rolling moraine hills surrounded by mountain ranges. The kettlehole is solely fed by rainwater, and frequently inundated over winter when potential evapotranspiration is low, but can also fill following a heavy rain event in summer. Within 50 meters from the edge of the kettlehole, there is a strong hydrological gradient from above the maximum flood line down to land submerged for an average of more than 250 days per year. Flooding data were collected using a capacitance probe that continuously recorded the water height above the lowest elevation in the kettlehole between November 2006 and March 2015.

##### 1.2 Plant community data

The kettlehole includes four replicated transects that run from near the lowest points in the wetland to above the flood line. Community composition was determined mid-February 2015, by identifying the presence and abundance of each individual plant to species within a total of 67 quadrats 25 × 25 cm in size. Quadrats were located at 2 m intervals along each transect beginning at 0 m (lowest point of kettlehole basin) and advancing upslope to the kettlehole margin. Foliar cover was estimated for each species using the following cover estimates: 0.5%, 1%, 2%, 3%, 4%, 5%, 10%, 15%, 20%, 30%, 40%, . . . , 100%. We restricted the analysis to the 15 most abundant species in the study area for which we sampled traits on at least 20 individuals.

The original study further included 120 supplementary quadrats that were not subjected to flooding and/or that were part of an experimental treatment (Purcell et al. 2019). Those quadrats were excluded from the present study but their functional trait data was used to increase the number of trait samples used to estimate species intraspecific trait distribution.

##### 1.3 Functional trait measurements

A single individual of each species present in each quadrat was located next to each quadrat for collection. The vegetative height (cm) of this specimen was measured before

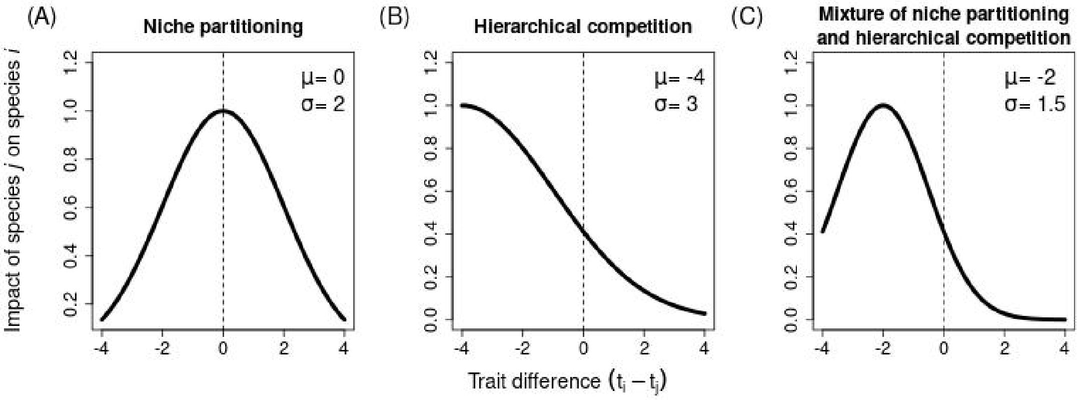


Supplementary Figure 1: Illustration of our modeling of interspecific pairwise interaction coefficients. They are defined as a function of trait differences that depend on peak position ($\mu$) and width ($\sigma$) parameters (see methods). (A) Pairwise interaction coefficients are higher for small trait differences, indicating a predominance of niche partitioning. (B) Species with a large trait value are competitively superior over species with small trait values. (C) Most of the interaction coefficients follow the same pattern as (A) but for small trait differences, hierarchical competition is the predominant process.

the whole plant was carefully removed, stored in a sealed ziplock bag with a moist paper towel within a chilled container. Depending on size, three, five, ten or 30 leaves were removed from each individual and weighed on analytical balances to obtain fresh mass (mg). Leaf area measurements were obtained from photographs of the leaves using ImageJ software. Dry masses of the leaf samples were obtained after oven drying samples at 60 °C. Dry weights and areas of leaf samples were used to calculate specific leaf area (SLA), the one-sided projected leaf area divided by dry mass (mm2 mg-1).

The root samples were transported to the University of Waikato where they were cleaned and tamped dry. The ‘microbalance method’ was used to measure an index of root porosity [8]. Ten short sections of fresh unsuberized roots, approximately 10 mm in length, were cut at least 10 mm back from the root tip. Whenever possible, sections were taken from first order terminal roots, otherwise second order roots with no secondary thickening were used. Roots were cleaned using fine brushes and surface moisture was removed using tissue paper prior to obtaining the initial mass (w1). The root sections were transferred into a 20 ml glass vial that was filled with water. The vial was placed into a vacuum desiccator three times for five minutes each, with pressure rapidly returned to atmospheric at the end of each five-minute period. The root sections were then removed from the vial, placed on dry tissue paper, and briefly rolled to remove surface moisture prior to obtaining the final mass (w2). Root porosity was calculated as $100 \times (w2 - w1)/w2$.

#### 1.4 Illustration of the pairwise interaction modeling framework

See Supplementary figure 1 and methods in the main article.

#### 1.5 Distribution of pairwise interaction coefficients

The average and standard deviation of pairwise interaction coefficients are essential features of interaction matrices in Lotka-Volterra models [1]. In our modeling approach, this is what differentiates the different biotic models: when the standard deviation of interspecific pairwise interaction coefficients approaches 0, the biotic model approaches the 'No traits' biotic model. Additionally, when the average of interspecific pairwise interaction coefficients approaches 0, then the interaction matrix approaches an identity matrix and thus resembles the abiotic model or the null model. It is thus important to establish the link between the model parameters and the average and standard deviation of pairwise interaction coefficients to better understand the properties of the model.

Here we study the average and standard deviation of the interspecific pairwise interaction coefficients $\alpha_{ij}$ in the univariate case (a single trait is used to estimate $\alpha_{ij}$). $\alpha_{ij}$ coefficients are function of the trait difference $\Delta t_{ij}$ between species $i$ and species $j$. The function $f$ is controlled by the parameters $C$, $\mu$ and $\sigma$.

As noted in the main text,

$$\alpha_{ij} = C \frac{1}{\sigma\sqrt{2\pi}} \exp\left[-\frac{1}{2}\left(\frac{t_k - t_i - \mu}{\sigma}\right)^2\right],$$

which can be written equivalently as:

$$\alpha_{ij} = C \times f(\Delta t_{ij}).$$

$f$ is a Gaussian distribution function of the trait difference $\Delta t_{ij} = t_i - t_j$ between species $i$ and $j$ and of parameters $\mu$ and $\sigma$. $C$ is a multiplicative constant.

The mean and variance of $\alpha_{ij}$ are:

$$E(\alpha_{ij}) = C \times \int_{-\infty}^{\infty} f(\Delta t_{ij}) g(\Delta t_{ij}) d\Delta t_{ij},$$

$$Var(\alpha_{ij}) = E(\alpha_{ij}^2) - E(\alpha_{ij})^2 = C^2 \times \int_{-\infty}^{\infty} f(\Delta t_{ij}) f(\Delta t_{ij}) g(\Delta t_{ij}) d\Delta t_{ij} - E(\alpha_{ij})^2,$$

and $g$ is the function that describes the distribution of interspecific trait differences $\Delta t_{ij}$. If there is an infinite number of species and if the functional trait distribution is Gaussian (i.e. normal and standardized), then $g$ is a Gaussian distribution function of mean 0 and standard deviation $\sqrt{2}$. Thus, both the mean and variance of $\alpha_{ij}$ depend on the integral of a product of Gaussian functions.

The product of Gaussian distribution functions (univariate or multivariate) is a Gaussian distribution function multiplied by a scaling coefficient [2]. Both the parameters of the function $N$ and the scaling coefficient $S$ can be derived analytically from the parameters of the initial Gaussian distribution functions [2].

In consequence,

$$f(x)g(x) = S_{fg} N_{fg}(x)$$

$$f(x)f(x)g(x) = S_{ffg} N_{ffg}(x)$$

with $N_{fg}(x)$ and $N_{ffg}(x)$ being Gaussian distribution functions and $S_{fg}$ and $S_{ffg}$ being scaling coefficients.

Then the mean and variance of $\alpha_{ij}$ are:

$$E(\alpha_{ij}) = C \times S_{fg} \int_{-\infty}^{\infty} N_{fg}(\Delta t_{ij}) \times d\Delta t_{ij}$$

$$Var(\alpha_{ij}) = C^2 \times S_{ffg} \int_{-\infty}^{\infty} N_{ffg}(\Delta t_{ij}) \times d\Delta t_{ij} - E(\alpha_{ij})^2$$

$N_{fg}$ and $N_{ffg}$ are Gaussian distribution functions so both of their integrals equal 1. This implies that

$$E(\alpha_{ij}) = C \times S_{fg}$$

$$Var(\alpha_{ij}) = C^2 \times (S_{ffg} - S_{fg}^2)\,.$$

The scaling coefficients $S_{ffg}$ and $S_{fg}$ depend only on $\mu$ and $\sigma$. The analytical expression of $S_{ffg}$ and $S_{fg}$ can be expressed using the demonstration of Bromiley (2003). A similar reasoning can be applied to the multivariate case.

The analytical forms of $E(\alpha_{ij})$ and $Var(\alpha_{ij})$ are not particularly intuitive. Thus to study their relationship to $\mu$ and $\sigma$, we coded the calculations of Bromiley (2003) and simulated the values of the average and standard deviation of interspecific pairwise coefficients as a function of parameters $C$, $\mu$ and $\sigma$ (Supplementary figure 2).

We also studied the bivariate case (Supplementary figure 3), where the average and standard deviation of interspecific pairwise coefficients are functions of $C$, $\mu_1$, $\mu_2$, $\sigma_1$, $\sigma_2$ and $\rho$. In that last case, we draw random values of $\sigma_1$, $\sigma_2$ and $\rho$. $\sigma_1$ and $\sigma_2$ were drawn from a uniform distribution between 0.05 and 10, while $\rho$ was drawn from a uniform distribution between -1 and 1. We then calculated the average and standard deviation of interspecific pairwise coefficients for three values of the pair ($\mu_1$, $\mu_2$) that represent (1) niche partitioning across both dimensions ($\mu_1 = 0$, $\mu_2 = 0$), (2) strong hierarchical competition across both dimensions ($\mu_1 = 3$, $\mu_2 = 3$) and (3) a mixed case ($\mu_1 = 0$, $\mu_2 = 3$).

In both the univariate and the bivariate case, we can see that there is a strong drop of the standard deviation of $\alpha_{ij}$ for values of $\sigma$, $\sigma_1$ and $\sigma_2$ that exceed 5.

#### 1.6 Prior definition

We specified the prior distributions of parameters so that they would be uninformative but still able to regularize the parameter space: typically we down-weighted the portions of the parameter space that were uninformative or where parameters become unidentifiable. Prior distributions are indicated in Supplementary Table 1.

- We set large, uninformative priors for $a$, $b$, $C$ and $\phi$.
- We constrained the $\mu$, $\mu_1$, $\mu_2$ with a prior distribution that mimic the distribution of trait differences. This was set so that the maximal value of $\alpha_{ij}$ (e.g. for $t_i - t_j = \mu$ in the single-trait case) would be found within the observed distribution of trait differences.
- We set a regularizing prior for $\sigma$ (single trait biotic model) and $\sigma_1$, $\sigma_2$ (multi-trait biotic model). As seen above, when those parameters reaches high values ($\sigma > 5$),

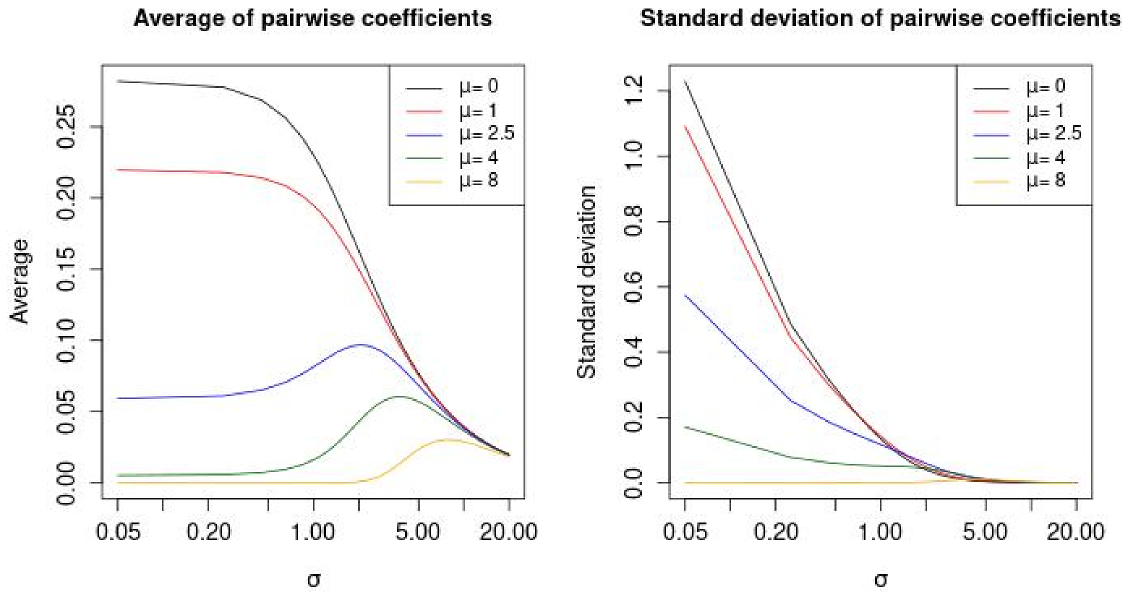


Supplementary Figure 2: Average (left) and standard deviation (right) of interspecific pairwise coefficients $\alpha_{ij}$ as a function of $\sigma$ and $\mu$ (C = 1). Note that the x-axis is on a log scale.

the variance of $\alpha_{ij}$ is close to 0. This represents a part of the parameter space where, in the univariate case:

$$\lim_{\sigma\to\infty} \alpha_{ij} = \lim_{\sigma\to\infty} \frac{C}{\sigma\sqrt{2\pi}} \exp\left[-\frac{1}{2}\frac{(t_i - t_j - \mu)^2}{\sigma^2}\right] \approx \frac{C}{\sigma\sqrt{2\pi}},$$

and in the multivariate case:

$$\lim_{\sigma\to\infty} \alpha_{ij} \approx \frac{C}{2\pi\sigma_1\sigma_2\sqrt{1-\rho^2}}$$

As such, the parameter $\mu$ (or parameters $\mu_1$ and $\mu_2$) becomes irrelevant and $C$ and $\sigma$ (or $C$, $\sigma_1$, $\sigma_2$, and $\rho$) are not identifiable. We thus set the priors for $\sigma$, $\sigma_1$, $\sigma_2$ with log-normal distributions that limit the sampling of large, uninformative values.

#### 1.7 Predicted community structure

For each assembly model, we estimated the predicted community diversity pattern, i.e. the predicted mean α-diversity and β-diversity and compared it to the observed diversity values and among assembly models. We estimated the mean α-diversity and β-diversity with and without including the spatial aggregation process modeled through the uncertainty parameter $\phi$ [3]. The mean α-diversity and β-diversity were estimated using the inverse of Simpson diversity index that consider species covers. We used the multiplicative partition of diversity ($\gamma = \beta \times \alpha$) to obtain independent α-diversity and β-diversity values [4].

Estimation without spatial aggregation (fixed): for each assembly models, α-diversity and β-diversity were computed from the site-by-species matrix predicted by Banquo,

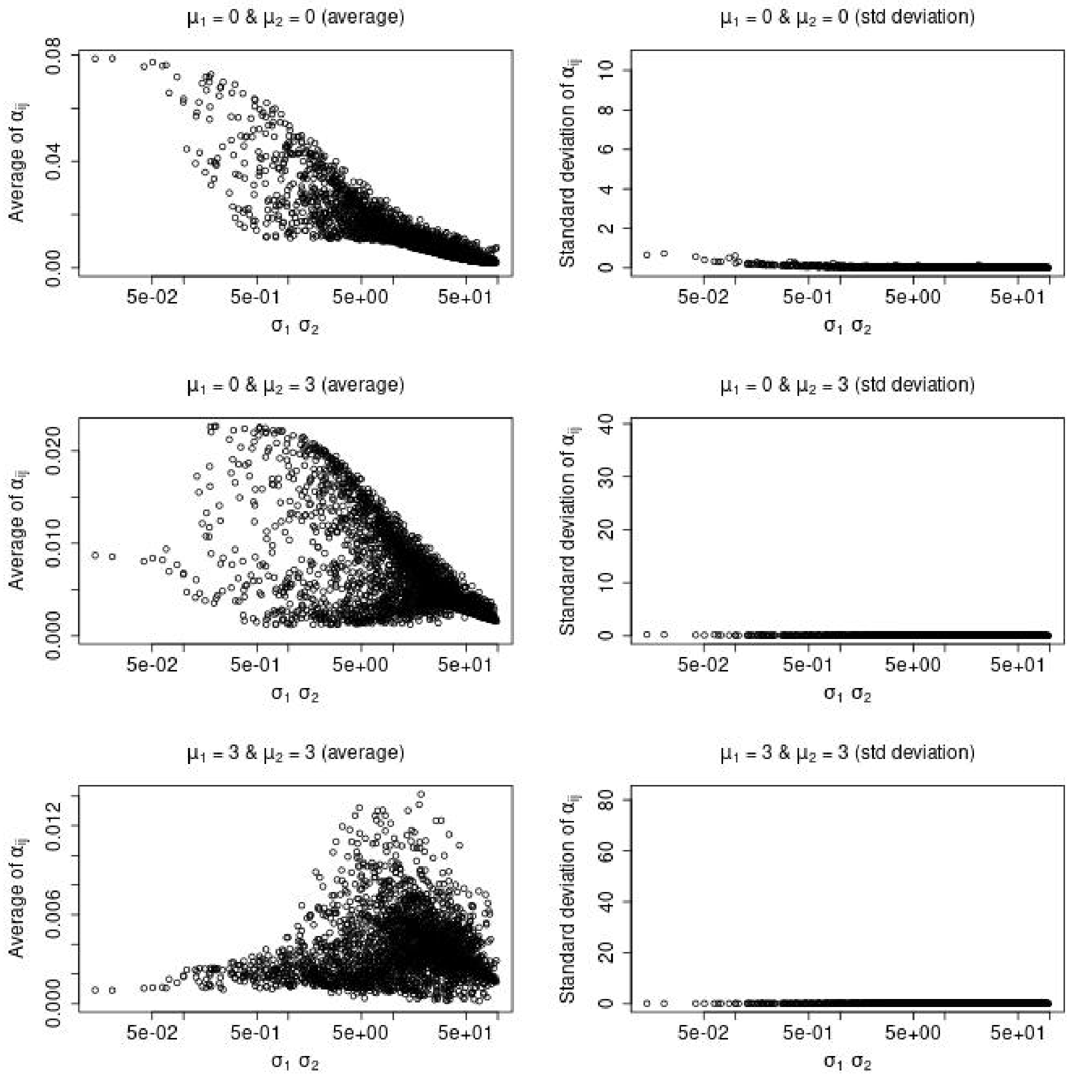


Supplementary Figure 3: Average (left column) and standard deviation (right column) of interspecific pairwise coefficients $\alpha_{ij}$ as a function of the product of $\sigma_1$ and $\sigma_2$. Each row represents a different set of values of $\mu_1$ and $\mu_2$. C is fixed at 1. Note that the x-axis is on a log scale.

| Parameter | Distribution | Bounds |
|---|---|---|
| All assembly models | | |
| $a$ | $U(0.01, 0.99)$ | $[0.01, 0.99]$ |
| $\phi$ | $Half - Cauchy(2.5)$ | $[0, \infty]$ |
| Abiotic model, Abiotic & Biotic models | | |
| $b$ | $U(0.025, 1.5)$ | $[0.025, 1.5]$ |
| Biotic models, Abiotic & Biotic models | | |
| $C$ | $logN(0.69, 2.3)$ | $[0.05, \infty]$ |
| Biotic models and abiotic & Biotic models One trait | | |
| $\mu$ | $N(0, 1.8)$ | $[-3.0, 3]$ |
| $\sigma$ | $logN(0, 3.5)$ | $[0.05, 10]$ |
| Biotic models and abiotic & Biotic models Two traits | | |
| $\mu_1$ | $N(0, 1.8)$ | $[-3.0, 3]$ |
| $\sigma_1$ | $logN(0, 2)$ | $[0.05, 10]]$ |
| $\mu_2$ | $N(0, 1.8)$ | $[-3.0, 3]$ |
| $\sigma_2$ | $logN(0, 2)$ | $[0.05, 10]$ |
| Biotic models and abiotic & Biotic models Two traits with interaction | | |
| $\rho$ | $N(0, 1)$ | $[-0.9, -0.9]$ |

Supplementary Table 1: Prior distribution of the assembly model parameters.

which contains the mean species cover along the flooding gradient. These fixed diversity estimates were computed across the parameter posterior. We anticipate that ignoring species spatial aggregation (i.e. species cover stochasticity) will provide a better comparison of the assembly models' outputs. We compared these values to the observed α-diversity and β-diversity of plant communities when species spatial aggregation along the flooding gradient is ignored. To do so, we estimated the α-diversity and β-diversity predicted by a stacked species distribution modelling approach [5] [7]: the cover of each individual was modeled as a function of the flooding gradient. We used generalized additive models and assumed that species cover was a beta-distributed variable using the R-package mgcv [9]. R-package mgcv does not implement the same likelihood that we used for Banquo (with cover classes), so as an approximation, species cover was instead assumed to be a continuous beta-distributed variable.

Estimation with spatial aggregation (stochastic): we first computed the site-by-species matrix predicted by Banquo at the median posterior. For each mean species cover value in each cell of the matrix, we then generated a random cover value drawn from a beta distribution with the calibrated uncertainty parameter φ. We then calculated mean α-diversity and β-diversity of this stochastic site-by-species matrix, and we repeated this procedure 500 times. We lastly compared the resulting stochastic diversity values with the observed α-diversity and β-diversity.

### 2 Supplementary results

#### 2.1 Posterior distributions

The complete prior and posterior distribution of all individual assembly models are displayed in Supplementary figure 7 (null model), Supplementary figure 8 (abiotic model), Supplementary figures 9, 10, 11, 12, 13 (biotic models) and Supplementary figures 14, 15, 16, 17, 18 (abiotic & biotic models).

#### 2.2 Receiver operating characteristic curves

The receiver operating characteristic (ROC) curve of the predicted sites-by-species cover matrix was computed for each calibrated assembly model. The sites-by-species cover matrix was estimated at the median of the posterior distribution. The ROC curves are displayed in Supplementary figure 6.

#### 2.3 Predicted community structure

Using the inverse of Simpson diversity index, the observed mean α-diversity of the wetland communities was 1.94 and the β-diversity was 5.28; when ignoring stochasticity along the flooding gradient (see methods), the mean α-diversity was 7.22 and the β-diversity was 1.33. The assembly models predicted different αand β-diversities along the flooding gradient 4 when ignoring stochasticity. All diversity values discussed in this paragraph refer to the median value across the posterior distribution of the assembly model. Trivially, the null model, predicted that the β-diversity would be equal to 1 and the α-diversity equal to 15 because in that model, all species have equal cover in every community. Similarly, the model with only biotic interactions did not predict any species turnover along the gradient (β= 1) but predicted a smaller α-diversity (between 12.6 and 15.0). The models that included abiotic filtering (with or without biotic interactions) were able to predict species turnover along the gradient with a β-diversity value of 1.09 for the abiotic filtering model and between 1.06 and 1.23 for the assembly models that also include biotic filtering. They also predicted a lower median α-diversity (11.9 without biotic interactions and [9.23-12.7] with biotic interactions). Overall, the assembly models closest to the observed diversity values when ignoring stochasticity (α-diversity: 7.22; β-diversity: 1.33) were the model that included abiotic filtering and estimated biotic interactions from SLA (median α-diversity = 9.23; median β-diversity = 1.23).

When we included plant cover stochasticity as calibrated by our modeling procedure, the simulated αand β-diversities were all close to the empirical observed values (Modeled α-diversity: [1.92, 1.97] vs. 1.94 for the observed α-diversity. Modeled β-diversity: [5.44, 7.11] vs. 5.28 for the observed β-diversity).

#### 2.4 Comparison of observed and modeled plant cover

Supplementary figure 5 compares the plant cover modeled by the abiotic model and the abiotic & biotic model (calibrated with Height and SLA without interactions) to the observed plant cover. The figure further compares the cover modeled by the two aforementioned assembly models, as well as their distribution.

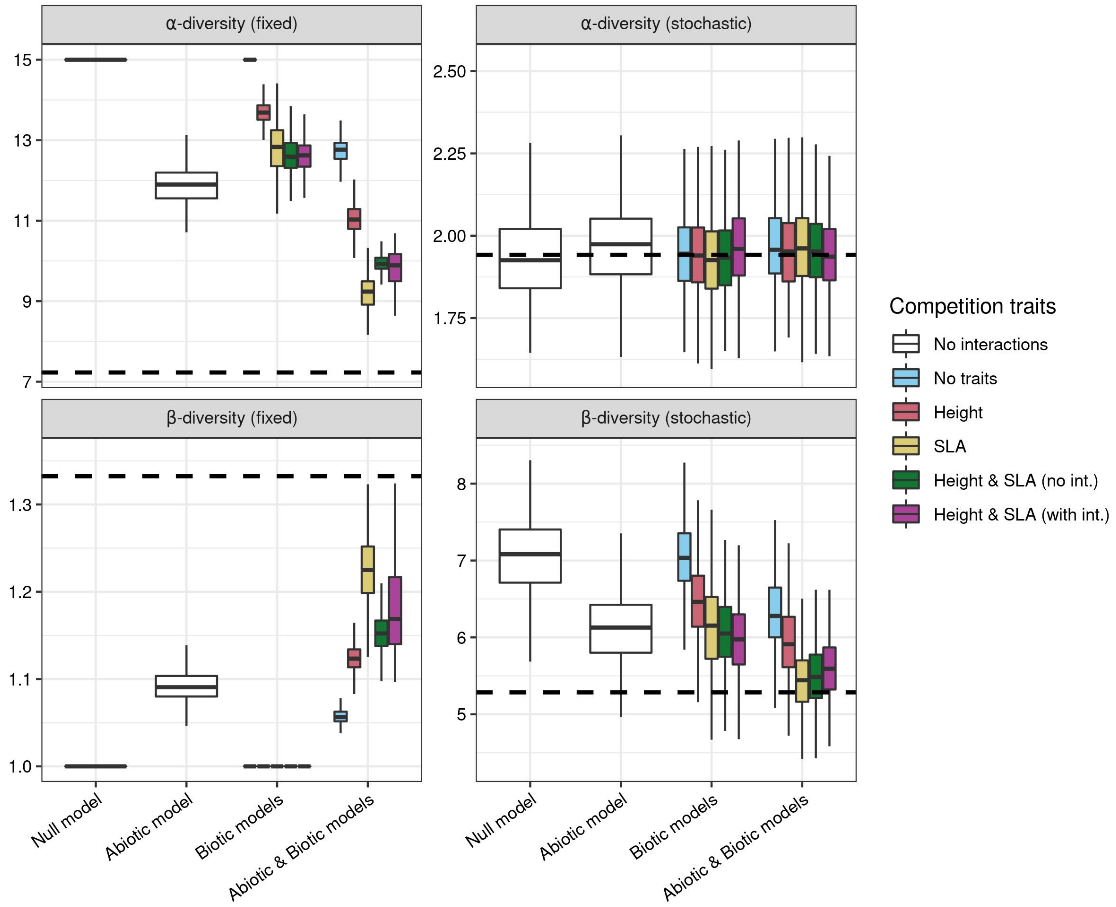


Supplementary Figure 4: αand β-diversities modeled by the assembly models. Assembly models are grouped by model type (Null, abiotic, biotic and abiotic & biotic). Boxplot color indicates the traits (or lack of thereof) used to compute the pairwise interaction matrix. The dotted horizontal line indicates the observed diversity value. In the left column are displayed the 'fixed' diversity estimates (without accounting for spatial aggregation). In the right column are displayed the 'stochastic' diversity estimates (accounting for spatial aggregation). See methods for more details.

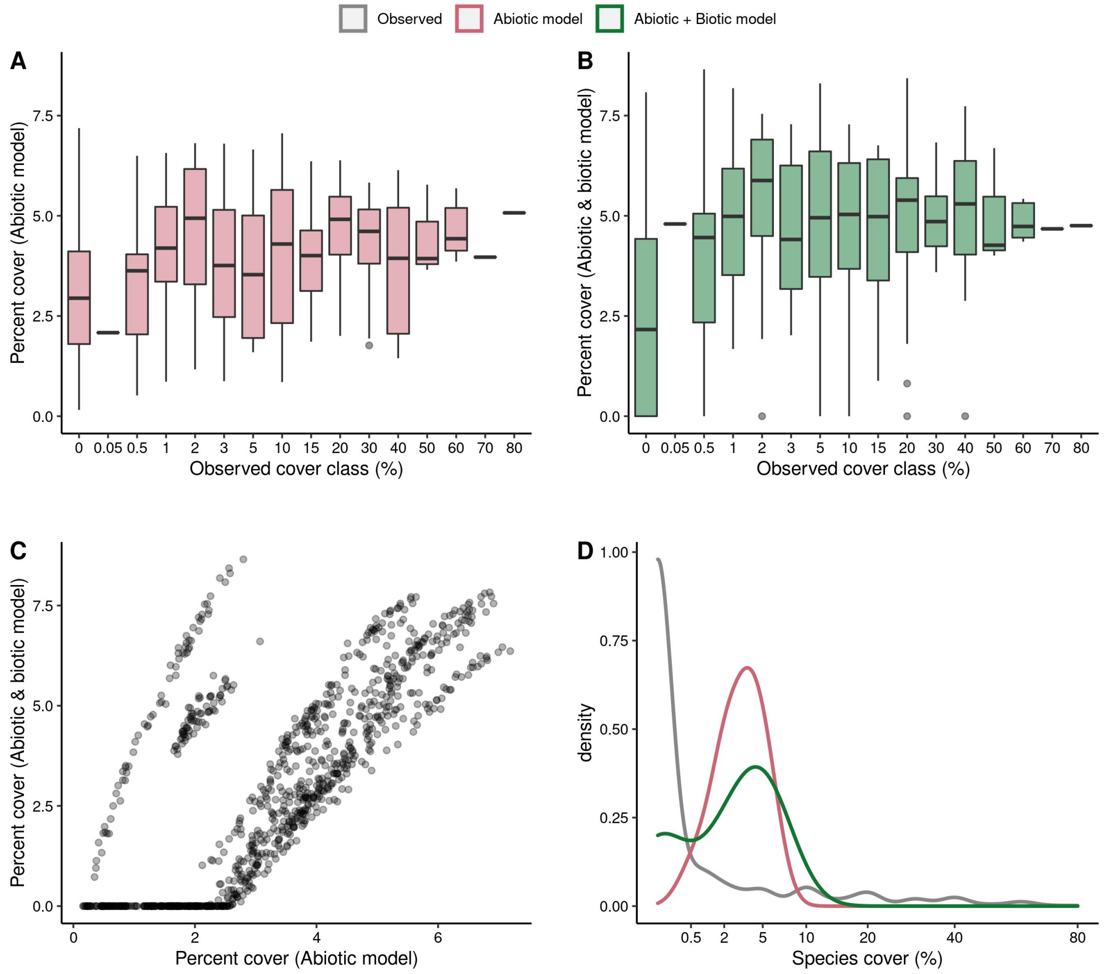


Supplementary Figure 5: Comparison of observed and modeled plant cover. Subpanels A and B show species cover modeled with abiotic filtering only (A), or abiotic filtering and SLA-based biotic interactions (B) as a function of observed plant cover classes. Subpanel C shows the species covers modeled with only abiotic filtering against the species covers modeled with both abiotic filtering and SLA-based biotic interactions. Subpanel D compares the density distribution of modeled species cover between those two models.

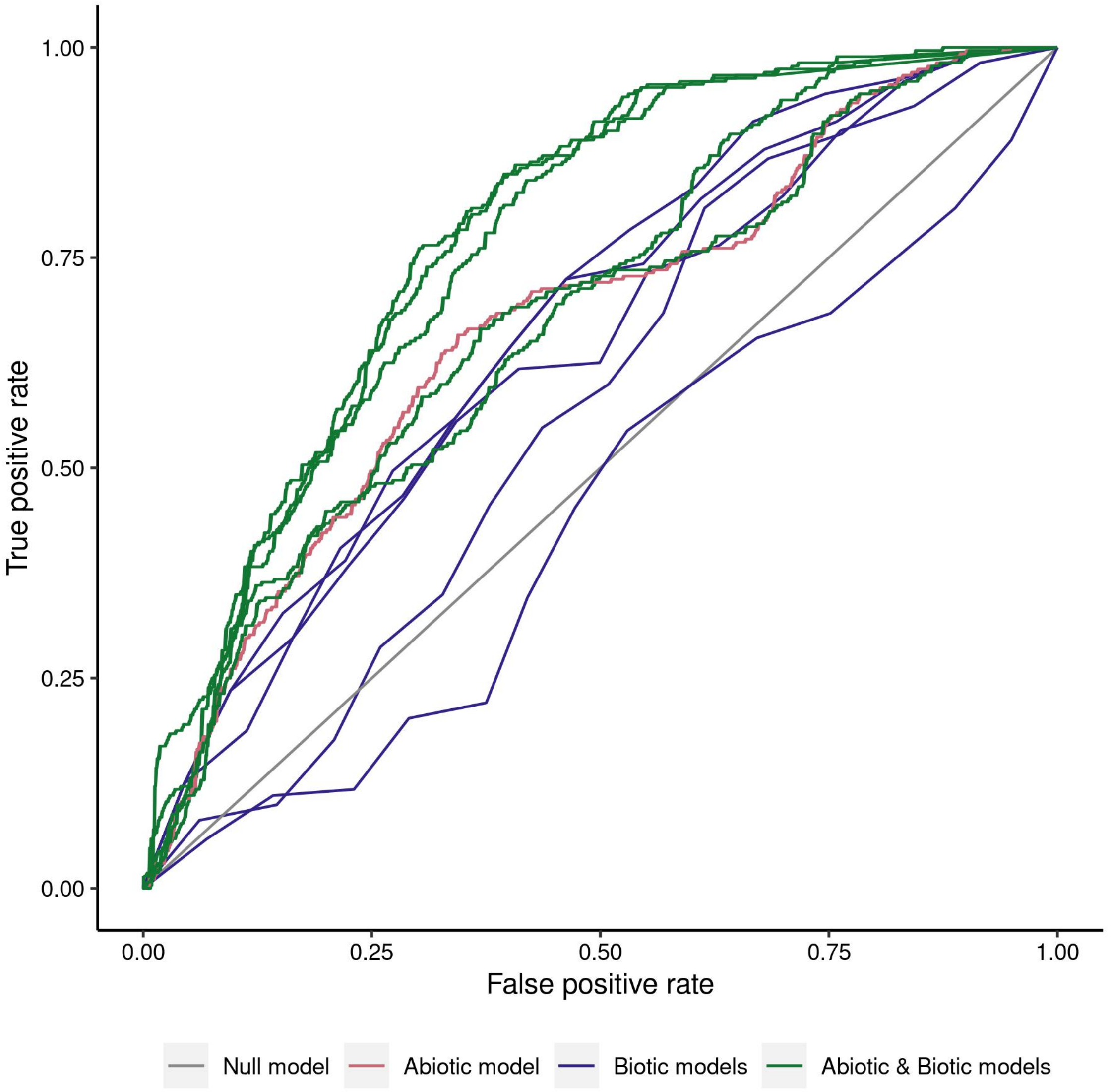


Supplementary Figure 6: ROC 'receiver operating characteristic' curves for the different assembly models: null model (grey), Abiotic model (pink), Biotic models (blue) and Abiotic & biotic models (green).

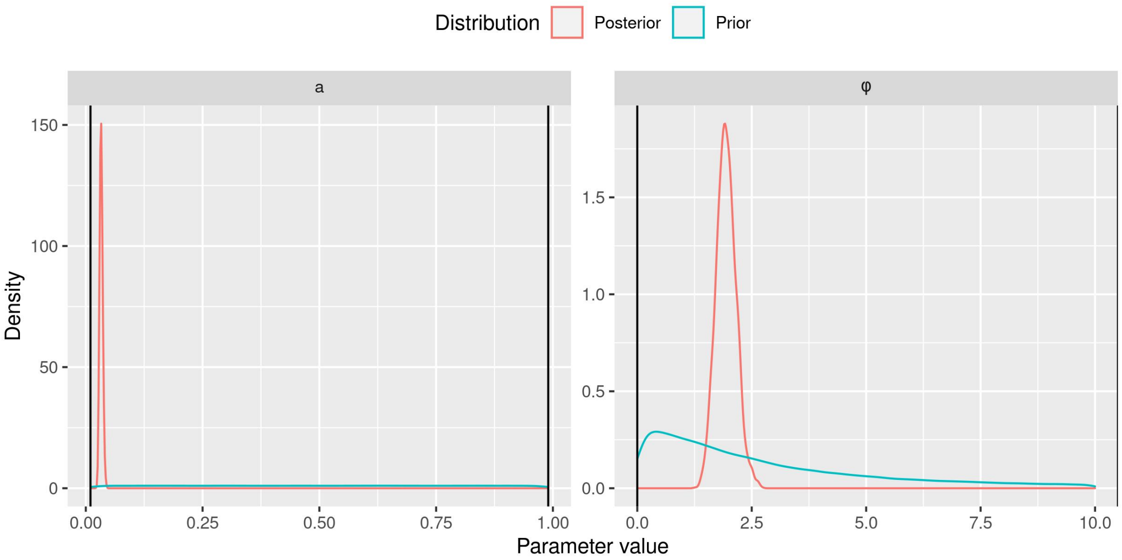


Supplementary Figure 7: Prior and posterior distribution of the parameters of assembly model 1 (Null model)

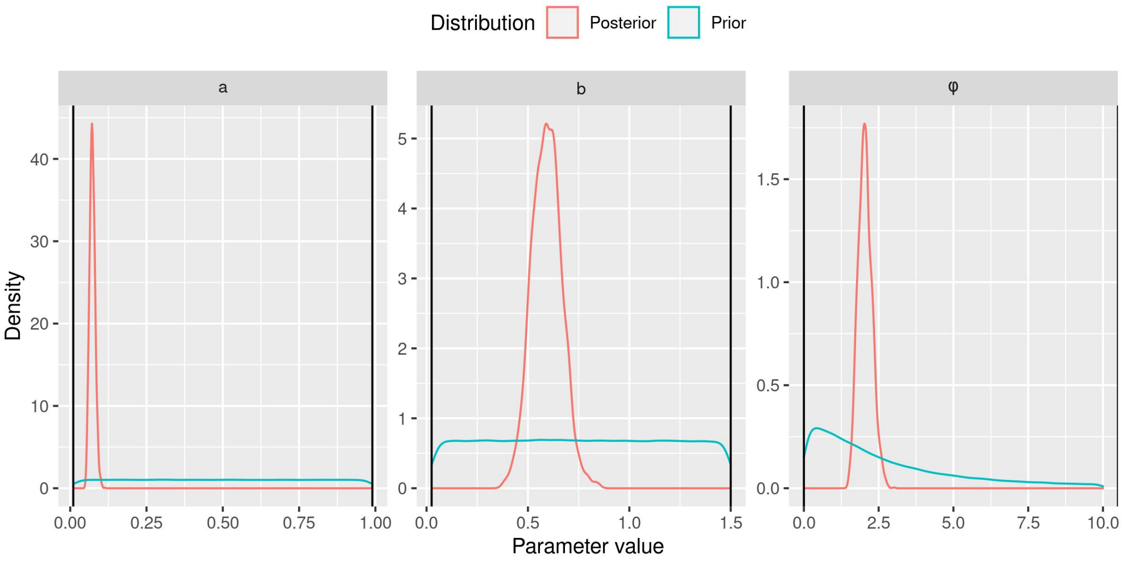


Supplementary Figure 8: Prior and posterior distribution of the parameters of assembly model 2 (Abiotic model)

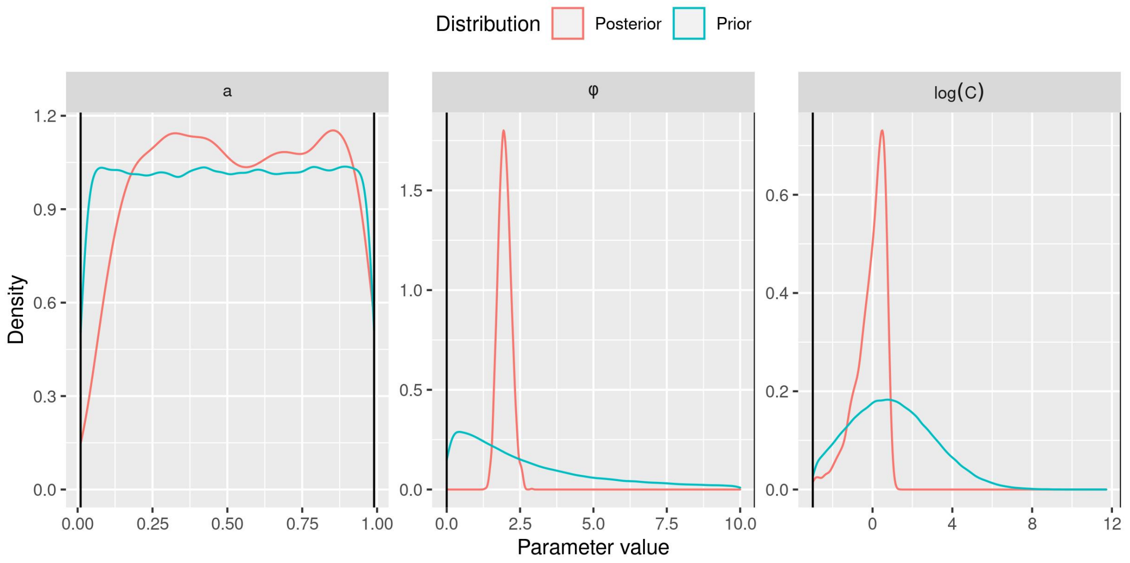


Supplementary Figure 9: Prior and posterior distribution of the parameters of assembly model 3 (Biotic model (no traits)). The parameter $C$ is displayed on a logarithmic scale.

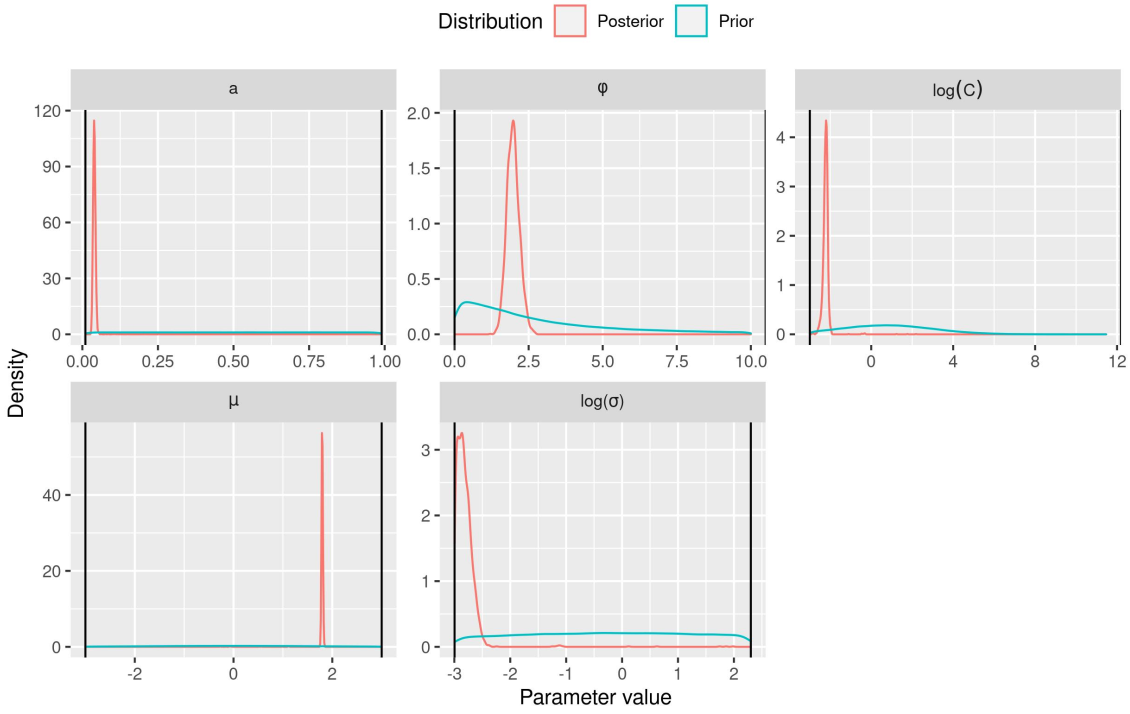


Supplementary Figure 10: Prior and posterior distribution of the parameters of assembly model 4 (Biotic model (Height)). The parameters $C$ and $\sigma$ are displayed on a logarithmic scale.

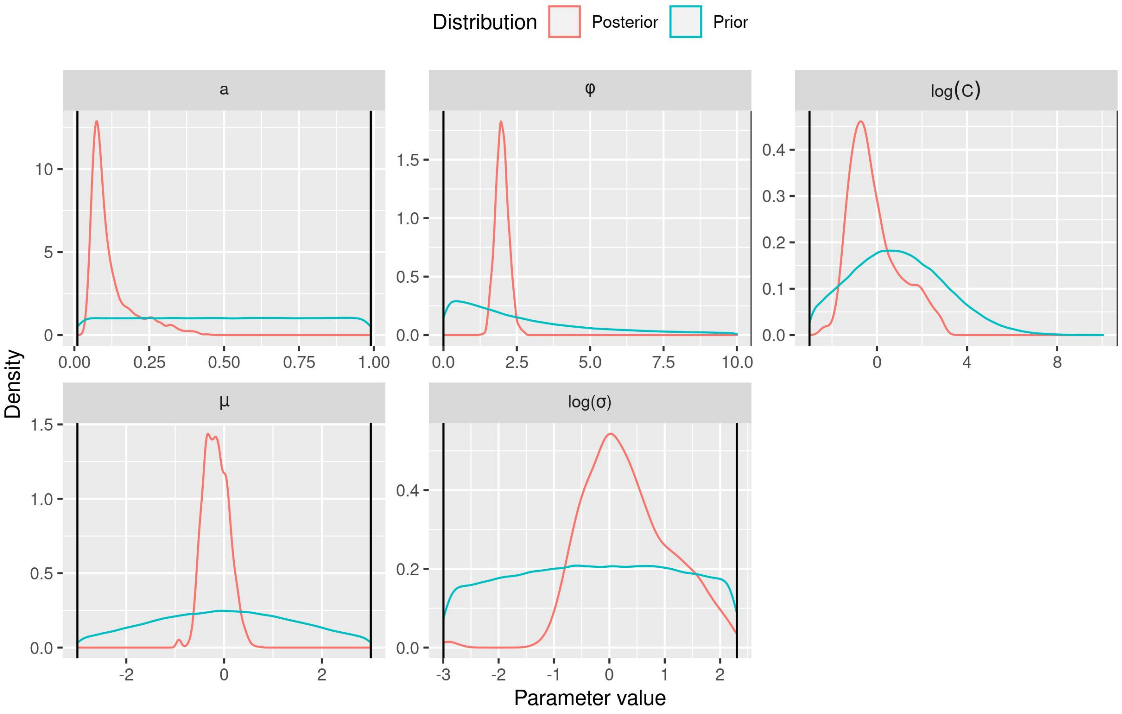


Supplementary Figure 11: Prior and posterior distribution of the parameters of assembly model 5 (Biotic model (SLA)). The parameters $C$ and $\sigma$ are displayed on a logarithmic scale.

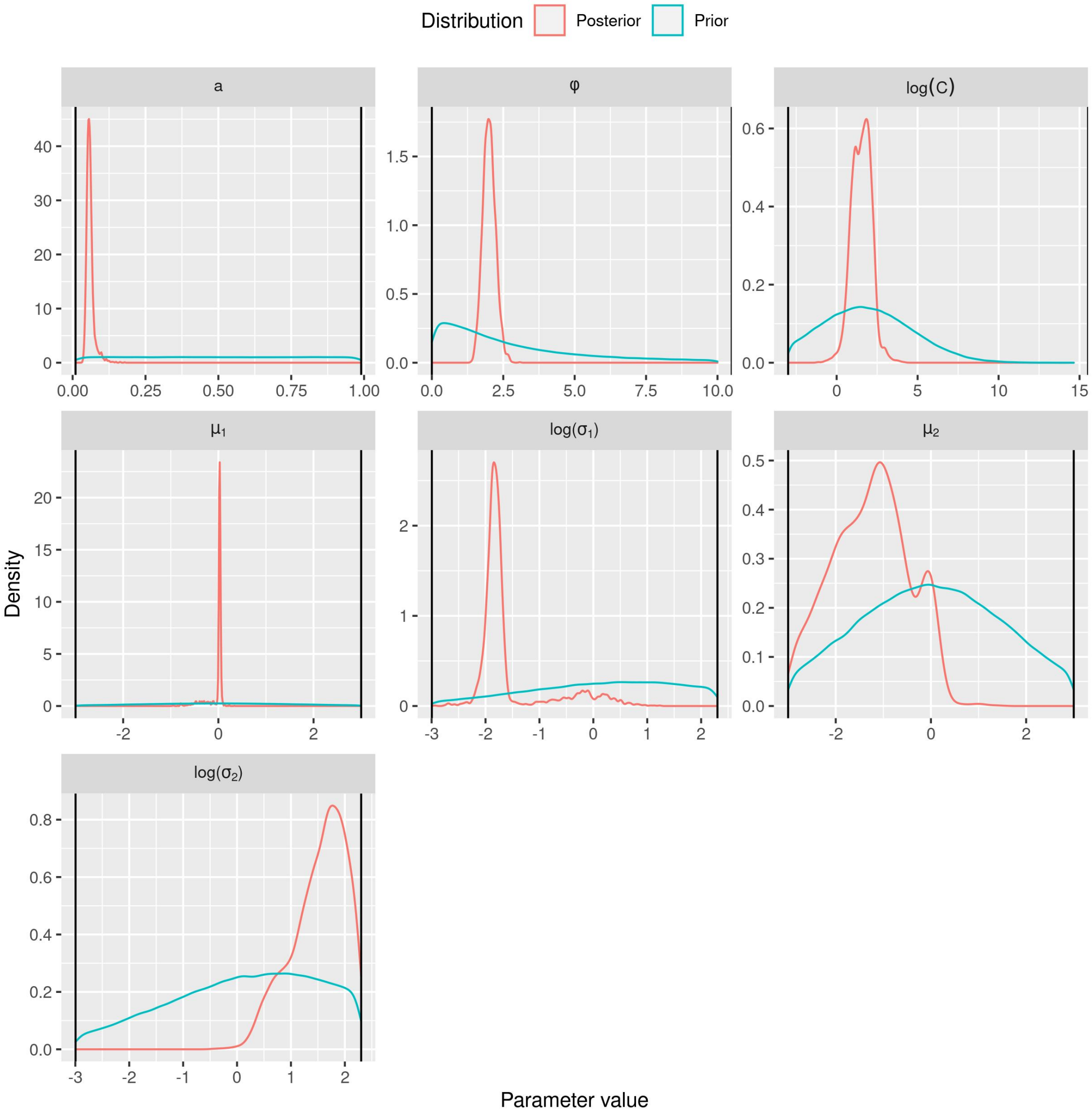


Supplementary Figure 12: Prior and posterior distribution of the parameters of assembly model 6 (Biotic model (Height & SLA, no interaction)). The parameters $C$, $\sigma_1$ and $\sigma_2$ are displayed on a logarithmic scale.

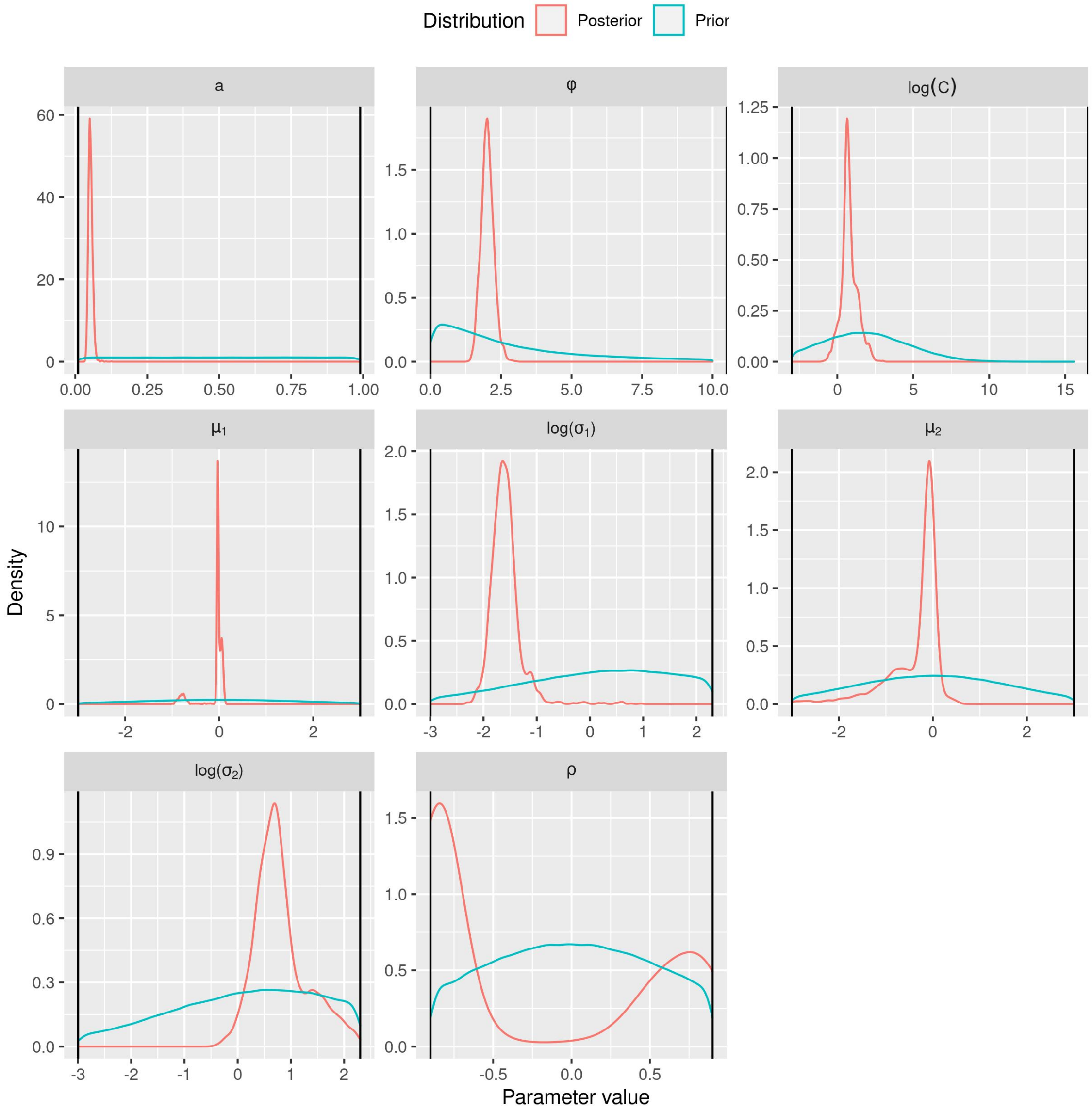


Supplementary Figure 13: Prior and posterior distribution of the parameters of assembly model 7 (Biotic model (Height & SLA, with interaction)). The parameters $C$, $\sigma_1$ and $\sigma_2$ are displayed on a logarithmic scale.

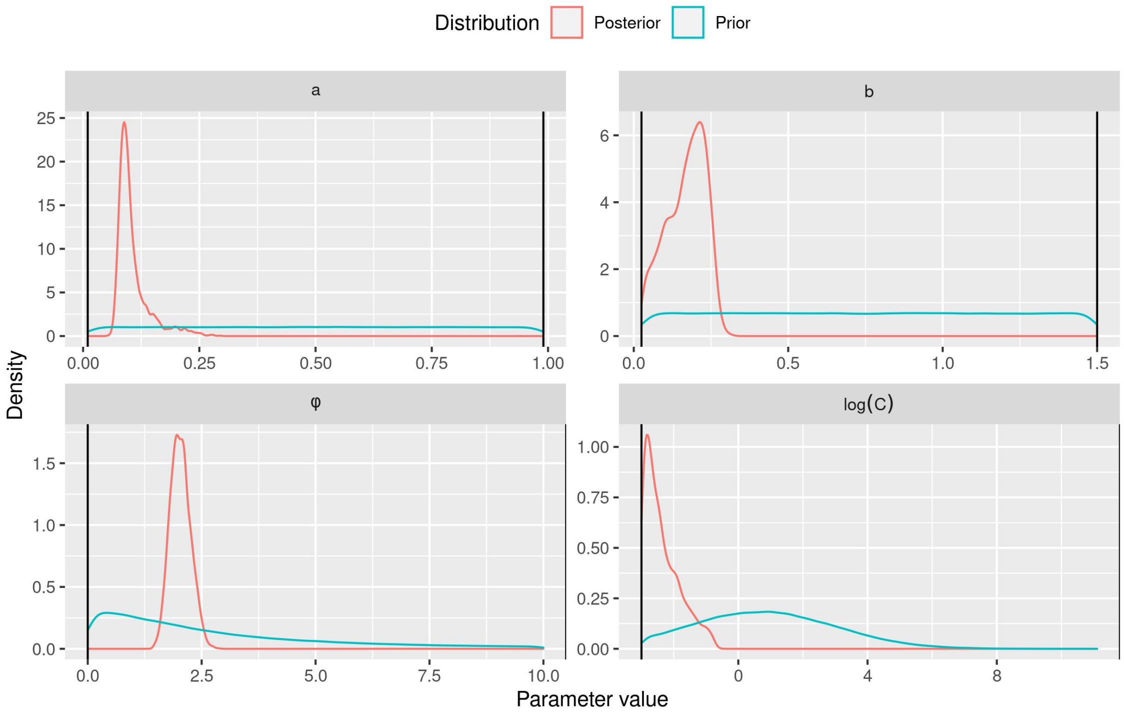


Supplementary Figure 14: Prior and posterior distribution of the parameters of assembly model 8 (Abiotic & Biotic model (no traits)). The parameter $C$ is displayed on a logarithmic scale.

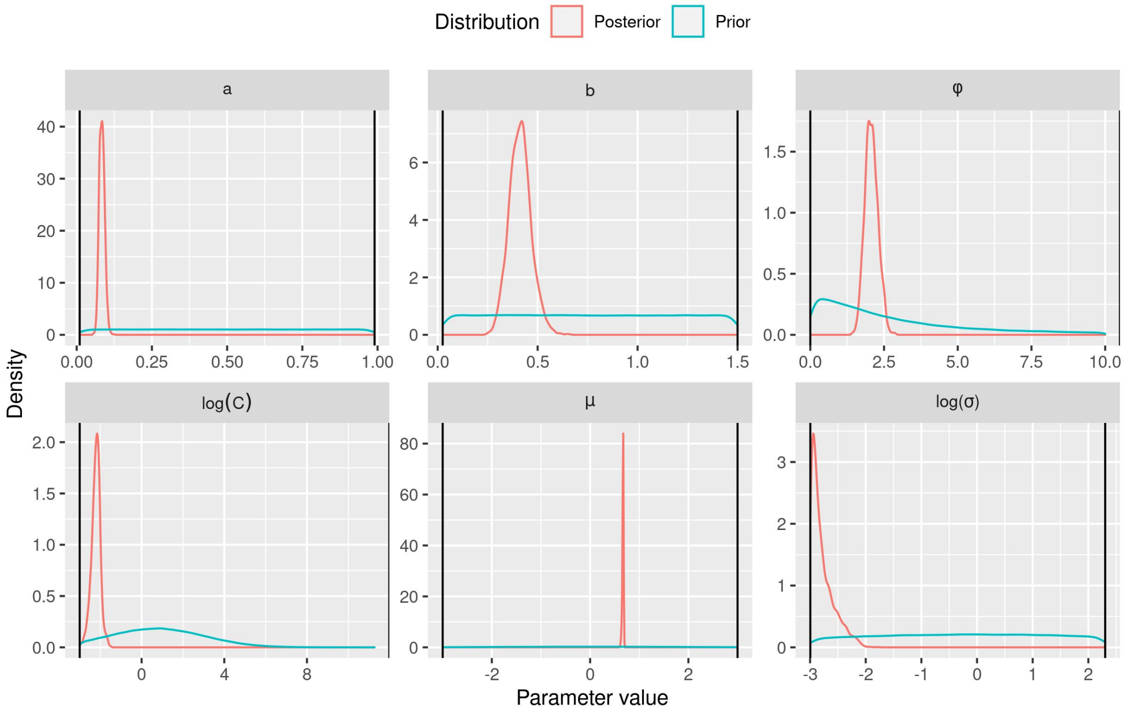


Supplementary Figure 15: Prior and posterior distribution of the parameters of assembly model 9 (Abiotic & Biotic model (Height)). The parameters $C$, $\sigma$ are displayed on a logarithmic scale.

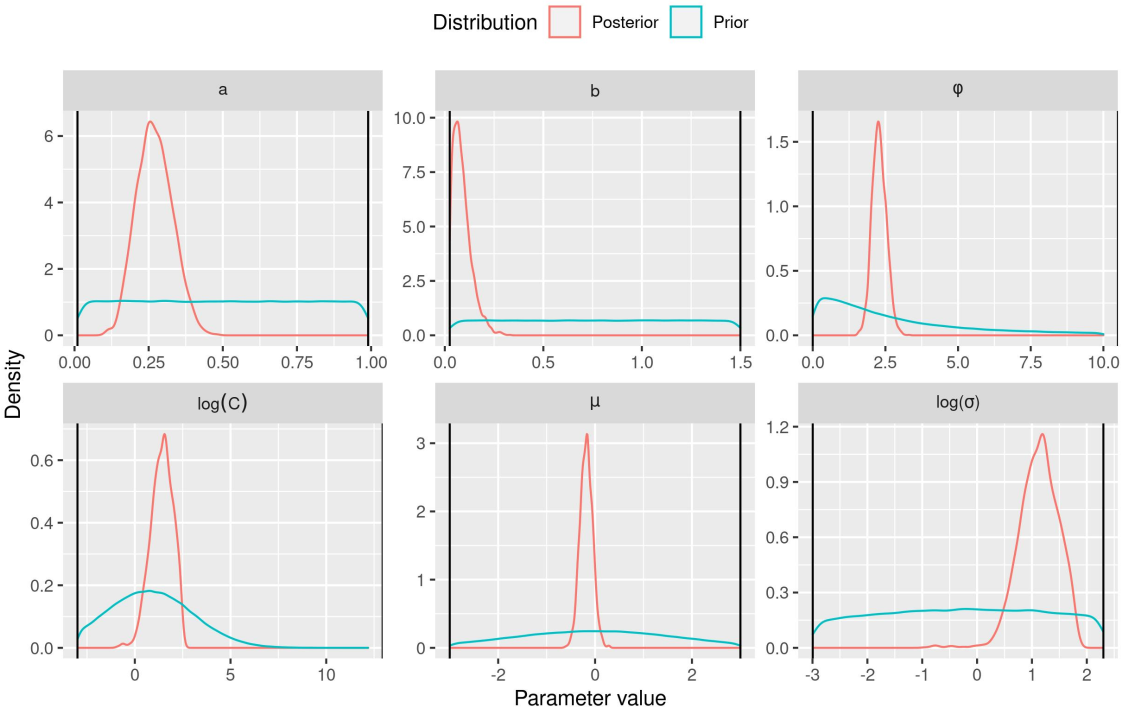


Supplementary Figure 16: Prior and posterior distribution of the parameters of assembly model 10 (Abiotic & Biotic model (SLA)). The parameters $C$, $\sigma$ are displayed on a logarithmic scale.

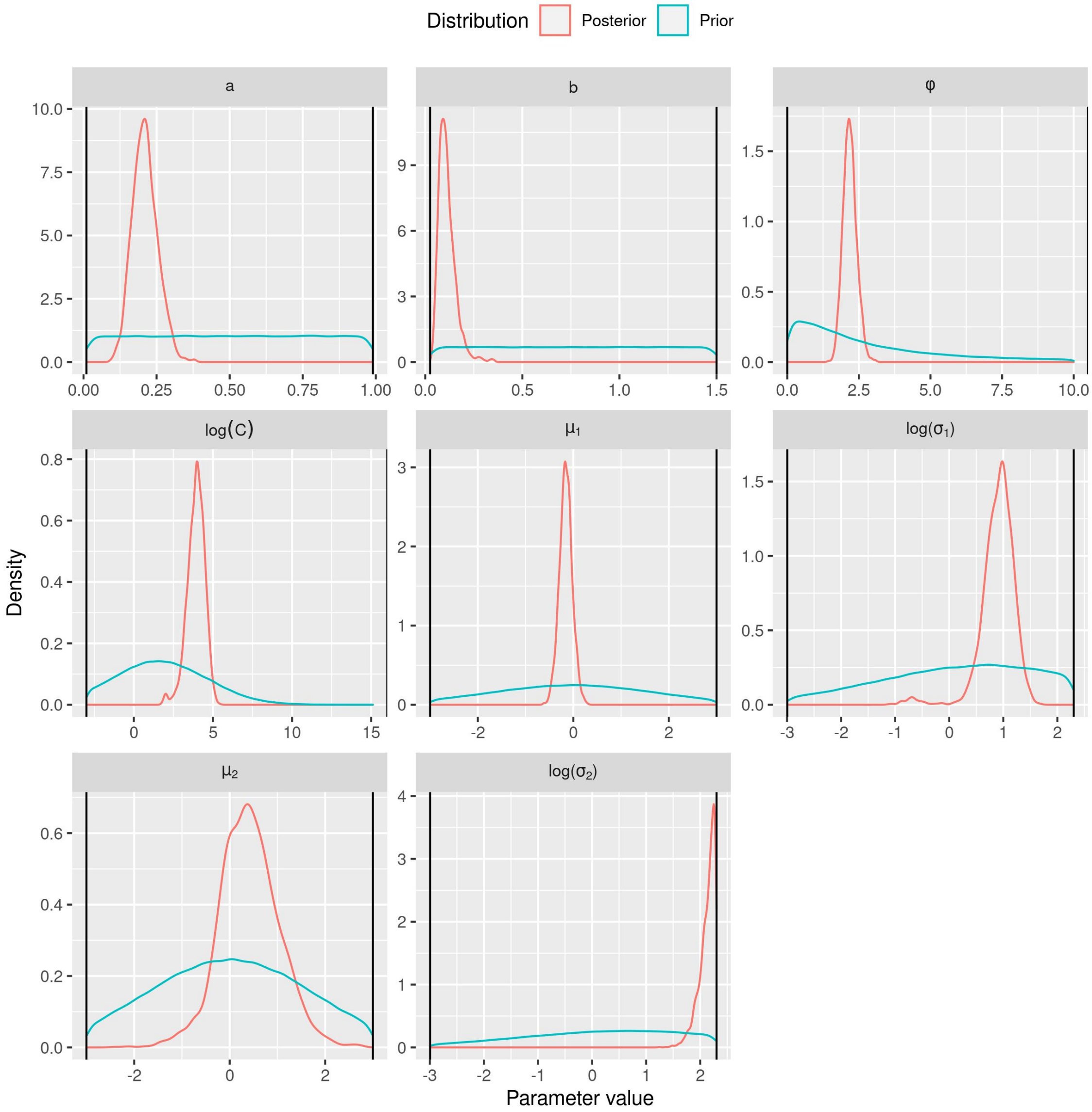


Supplementary Figure 17: Prior and posterior distribution of the parameters of assembly model 11 (Abiotic & Biotic model (Height & SLA, no interaction)). The parameters $C$, $\sigma_1$ and $\sigma_2$ are displayed on a logarithmic scale.

Supplementary Figure 18: Prior and posterior distribution of the parameters of assembly model 12 (Abiotic & Biotic model (Height & SLA, with interaction)). The parameters $C$, $\sigma_1$ and $\sigma_2$ are displayed on a logarithmic scale.
